# RosettaDDGPrediction for high-throughput mutational scans: from stability to binding

**DOI:** 10.1101/2022.09.02.506350

**Authors:** Valentina Sora, Adrian Otamendi Laspiur, Kristine Degn, Matteo Arnaudi, Mattia Utichi, Ludovica Beltrame, Dayana De Menezes, Matteo Orlandi, Olga Rigina, Peter Wad Sackett, Karin Wadt, Kjeld Schmiegelow, Matteo Tiberti, Elena Papaleo

**Affiliations:** Cancer Structural Biology, Danish Cancer Society Research Center, 2100, Copenhagen, Denmark; Cancer Systems Biology, Section for Bioinformatics, Department of Health and Technology, Technical University of Denmark, 2800, Lyngby, Denmark; Department of Clinical Genetics, Copenhagen University Hospital Rigshospitalet, Denmark; Department of Pediatrics and Adolescent Medicine, University Hospital Rigshospitalet, Copenhagen, Denmark; Institute of Clinical Medicine, Faculty of Medicine, University of Copenhagen, Copenhagen, Denmark

**Keywords:** Free energy calculations, binding free energy, folding free energy, Rosetta

## Abstract

Reliable prediction of free energy changes upon amino acidic substitutions (ΔΔGs) is crucial to investigate their impact on protein stability and protein-protein interaction. Moreover, advances in experimental mutational scans allow high-throughput studies thanks to sophisticated multiplex techniques. On the other hand, genomics initiatives provide a large amount of data on disease-related variants that can benefit from analyses with structure-based methods. Therefore, the computational field should keep the same pace and provide new tools for fast and accurate high-throughput calculations of ΔΔGs. In this context, the Rosetta modeling suite implements effective approaches to predict the change in the folding free energy in a protein monomer upon amino acid substitutions and calculate the changes in binding free energy in protein complexes. Their application can be challenging to users without extensive experience with Rosetta. Furthermore, Rosetta protocols for ΔΔG prediction are designed considering one variant at a time, making the setup of high-throughput screenings cumbersome. For these reasons, we devised RosettaDDGPrediction, a customizable Python wrapper designed to run free energy calculations on a set of amino acid substitutions using Rosetta protocols with little intervention from the user. RosettaDDGPrediction assists with checking whether the runs are completed successfully aggregates raw data for multiple variants, and generates publication-ready graphics. We showed the potential of the tool in selected case studies, including variants of unknown significance found in children who developed cancer, proteins with known experimental unfolding ΔΔGs values, interactions between target proteins and a disordered functional motif, and phospho-mimetic variants. RosettaDDGPrediction is available, free of charge and under GNU General Public License v3.0, at https://github.com/ELELAB/RosettaDDGPrediction.

## Introduction

Predicting the impact of amino acid substitutions in a protein or at a protein-protein interface is becoming more and more relevant as high-throughput sequencing data reveal a high rate of sequence polymorphisms of unknown functional significance in protein-coding regions^1^. In this context, multiplex-based assays provide a massive amount of data that can be complemented by structural studies on the effects of protein variants ^2–6^. Furthermore, saturation mutagenesis is experimentally very accessible thanks to the advances in multiplex technologies.

Therefore, molecular modeling approaches must keep the same pace and continue developing toward highthroughput applications.

A convenient and quantitative manner for assessing the impact of amino acid substitutions related to coding variants is based on estimating the changes in Gibbs free energy of folding/unfolding or binding. In this context, several computational approaches based on the analysis of protein structures are available to predict free energy changes upon mutation (ΔΔGs) in protein structures ^7–16^. These measurements can be used to classify the effect of disease-related variants on protein structural stability and, consequently, alterations of the cellular level or propensity to aggregation or proteasomal degradation ^17,18^, along with functional effects due to local changes in the interactions with other proteins or biomolecules ^19–21^.

Rosetta provides a variety of protocols to estimate changes in free energy in terms of binding and folding/unfolding ^8,11,12,16,22^. Most of these protocols estimate the change in free energy as an average over the free energy changes calculated in an ensemble of paired wild-type/mutated structures.

Rosetta protocols for the prediction of free energy changes upon mutation are characterized by three features: (i) the sampling method employed to generate the structural ensemble, (ii) the energy function used to quantify the free energy associated with each structure, and (iii) the degree of flexibility allowed in the structure to accommodate the mutation. Currently, three state-of-the-art strategies are available in Rosetta to estimate the change in either folding or binding free energy upon mutation. The first one, presented by Park and coworkers ^16^ and referred to as *cartddg*, is designed to work on monomeric proteins. It depends on sampling in Cartesian space (as opposed to internal dihedrals sampling), the *ref2015* Rosetta energy function (Cartesian space version), and small local backbone movements allowed in a three-residue window around the mutation site, together with side-chains movements within a 6 Å radius from the mutation site. The second protocol, *cartddg2020*, represents an updated variant of *cartddg* ^8^.

The third protocol, developed by Barlow and coworkers^11^ and named here *flexddg*, deals with estimating the changes in binding free energy upon mutation in a protein complex. It uses the “backrub” sampling method ^13^, which aims to recapitulate local backbone motions observed in crystal lattices. The *flexddg* protocol seems to perform better with the *talaris2014* energy ^11^ function. This protocol for binding free energies relies on a local sampling of backbone and side chains for residues within an 8 Å radius from the mutation, followed by global optimization of the side chains

Rosetta is a feature-rich software suite under active development, backed by a sizable community of users, and built over roughly 20 years. Running these protocols directly with Rosetta requires an extensive computational background and prior exposure to several Rosetta features. These requirements may discourage users with a more biology-oriented skillset, despite the benefit that accurate predictions of free energy changes upon mutations may bring to their research. Furthermore, Rosetta protocols for ΔΔG prediction are designed to be run considering one mutation at a time exclusively, making high-throughput screenings cumbersome to set up. We recently faced a similar challenge with implementing high-throughput scans based on the FoldX free energy function and making them parallelizable, more easily approachable, and applicable to structural ensembles. This led to the development of MutateX ^23^. FoldX, however, is known to suffer from limitations due to backbone stiffness during the sampling ^24^ and often low accuracy in predicting mutations with stabilizing effects on stability, even though most prediction methods are biased towards destabilizing mutations ^24,25^. Rosetta-based calculations could offer a valuable complement to the ΔΔG estimates currently accessible with MutateX. Thus, we developed RosettaDDGPrediction, a Python wrapper to perform Rosetta-based protocols for ΔΔG prediction. RosettaDDGPrediction’s outputs can also be converted to a format compatible with the MutateX plotting system, allowing for an expanded visualization toolkit. Here, we illustrate the applications and limits of the approach to four different cases of study, covering both methodological and biological applications. At first, we focused on the comparison with experimentally determined unfolding ΔΔG values (Case Study 1) and the influence of using AlphaFold2 models as starting structures (Case Study 2). Then, we showed two examples of applications of biological interest to study protein-protein interactions and post-translational modifications (Case Study 3) and to assess the functional impact of mutations identified by whole genome sequencing to address cancer predisposition (Case Study 4).

## Results

### Overview of the package

RosettaDDGPrediction is a pure Python package providing a uniform and easily accessible command-line interface to *flexddg, cartddg*, and *cartddg2020* protocols for the calculation of free energy changes upon mutation. It is devised to help users unfamiliar with the Rosetta suite perform mutational scans and collect, aggregate, and visualize data from those scans in an intuitive fashion. In RosettaDDGPrediction, a “protocol” is intended as a set of Rosetta runs and Python-based processing steps, which takes as inputs the three-dimensional structure of the protein of interest and a list of mutations to be performed, finally returning the predicted free energy changes associated with each input mutation. The *flexddg* protocol consists of only one call to the *rosetta_scripts* executable for each mutation, which performs all the necessary calculations as defined by Barlow and coworkers ^11^. On the other hand, the *cartddg* protocol first energetically relaxes the input structure by using the Rosetta *relax* program to generate an ensemble of relaxed conformations, followed by the selection of the most suitable one. Finally, it uses the *cartesian_ddg* application to relax the structure further and perform the free energy calculations. *cartddg2020* protocols represent updated versions of the original *cartddg* protocols. Here, the relaxation is performed by a Rosetta script passed to the *rosetta_scripts* executable, and then *cartesian_ddg* is run on the lowest energy structure produced by the relaxation. It is worth noting that the relaxation procedure produces only one structure, as per the original files provided with the work first describing the *cartddg2020* protocol ^8^. However, if the user decides to produce several relaxed structures, the most suitable one (according to user-selected criteria) will then be passed to *cartesian_ddg*. The standard protocols are described in specific YAML files provided with the package. With these files, expert users can still tap into the full potential of the Rosetta interface by providing virtually any Rosetta-compatible option to the executables used by each protocol.

RosettaDDGPrediction consists of four main executables (*rosetta_ddg_run, rosetta_ddg_check_run, rosetta_ddg_aggregate, rosetta_ddg_plot*) performing different tasks (**Figure 1**). Their behavior is controlled by a set of configuration files, which can be fully customized to fine-tune the parameters of each protocol, aggregation options, and plot aesthetics.

**Figure 1.**
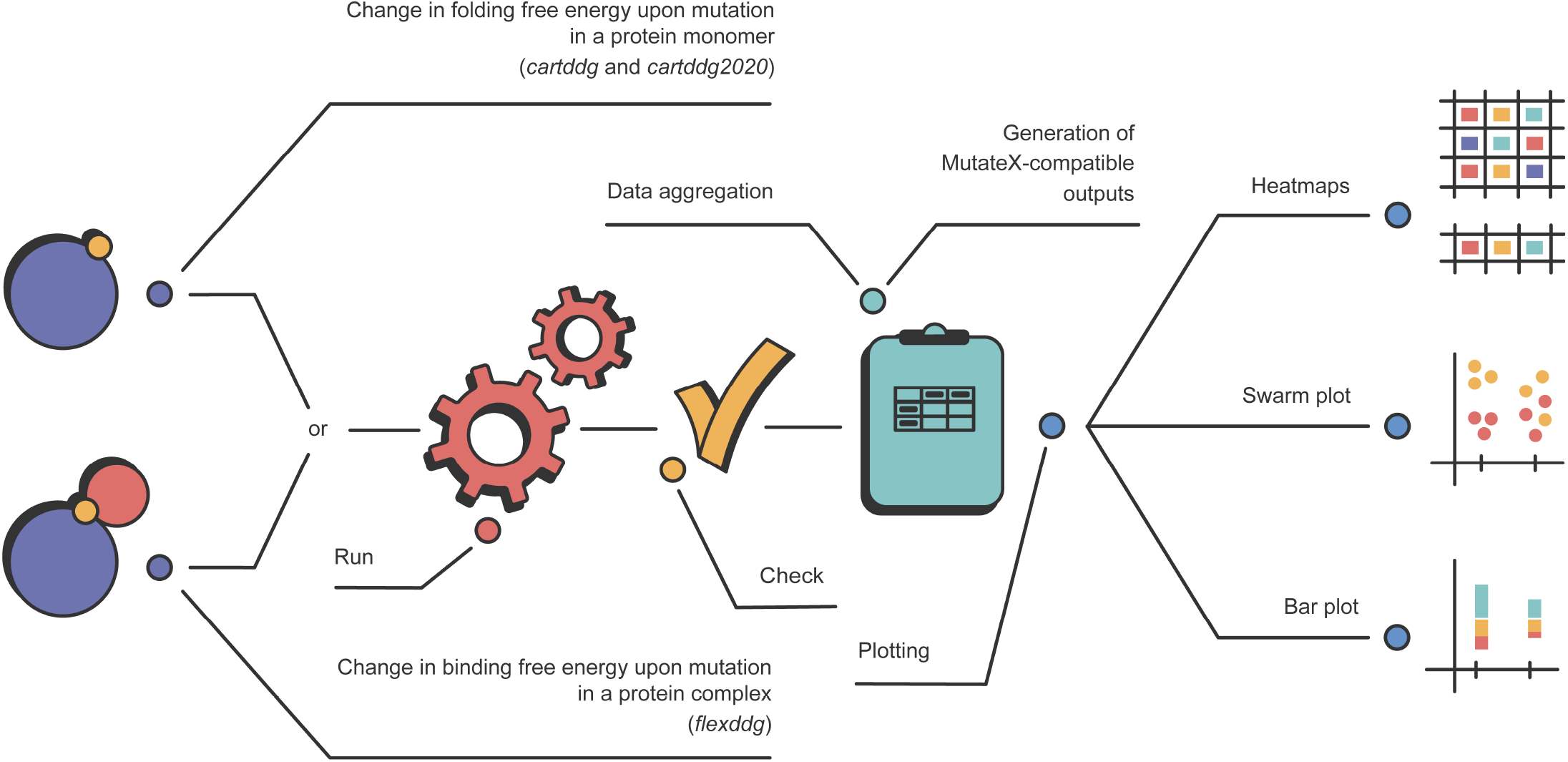
The RosettaDDGPrediction workflow and schematized plot types. The first step consists in running the *rosetta_ddg_run* executable to obtain the predicted ΔΔG values for the changes in folding free energy (for monomeric proteins) or binding free energy (for protein complexes). Then, *rosetta_ddg_check* can be used to ensure that all runs have been completed successfully. Data aggregation can then be performed with *rosetta_ddg_aggregate*, and aggregate data can finally be visualized in different ways (heatmaps, bar plots, swarm plots) using *rosetta_ddg_plot*.

*rosetta_ddg_run* is the executable responsible for running a Rosetta protocol to predict free energy changes upon mutation over a set of selected mutations. Given a protein structure in PDB format and a set of mutations, it generates all the data structures and configuration files to perform several runs in parallel, making them straightforward to perform and making the most of modern many-cores computing infrastructures. *rosetta_ddg_run* can optimize the workload distribution over the available resources to ensure efficient scheduling of the runs, thanks to the Dask Python package operating under the hood. *rosetta_ddg_run* easily handles multi-step protocols, requiring sequential Rosetta calls and possibly processing the output data between the steps. For example, for the aforementioned *cartddg* and *cartddg2020* protocols, *rosetta_ddg_run* takes care of both Rosetta calls and the processing steps.

Once the runs are completed, users can perform a sanity check on the calculations using *rosetta_ddg_check_run*, which identifies problematic runs by scraping the Rosetta output files. If the runs have been completed successfully, *rosetta_ddg_aggregate* can aggregate raw data from the large numbers of collected mutation runs into easily readable table files. These aggregate files contain, together with the calculated differences in free energy, additional information about each mutation, the Rosetta energy function used, and the number of structures generated in the final ensemble of structures. *rosetta_ddg_aggregate* also allows generating aggregate outputs compatible with the MutateX plotting system. Indeed, MutateX offers additional visualization tools, including density plots, logo plots, distribution plots, and summary tables that can be easily navigated ^23^.

Finally, *rosetta_ddg_plot* provides plotting utilities to explore the aggregated data through several visualization types, such as one-dimensional or two-dimensional heatmaps. The latter is particularly convenient when a saturation mutagenesis scan is run on a set of positions. The contribution of each term of the energy function to the final ΔΔG values may be visualized as a stacked bar plot, where positive and negative contributions add up on the corresponding semiaxes. Finally, since all protocols implemented so far in RosettaDDGPrediction determine the ΔΔG value associated with a mutation by averaging over the values produced by an ensemble of structures, the user may want to visualize the distribution of such values to investigate the source of potential outliers that may bias the average. In this case, a swarm plot displaying such values as separate data points is a very insightful overview provided by *rosetta_ddg_plot*.

To guide the user on the number of cores and time required for calculation, depending on the RosettaDDGPrediction protocol, energy function, and protein size, we report the results for different saturation scans in **Table 1**.

**Table 1.** Examples of performances of RosettaDDGPrediction for different protein sizes, the number of cores, and protocols applied. In the case of complexes the ‘Protein size’ column includes two values, i.e., one value for each protein/peptide in the complex. The one marked with a * is the one for which the saturation mutational scan was carried out. Calculations were run on servers equipped with either dual Xeon 6142 processors or dual Xeon 6242 processors. Each processor features 32 cores. The estimate refers to calculations done with Rosetta version 3.12.

| Protein size | Number of cores | Protocol | Time (hours) |
| --- | --- | --- | --- |
| 120 | 16 | cartddg (ref2015) | 33 |
| 250 | 24 | cartddg (ref2015) | 67 |
| 250 | 8 | cartddg (talaris2014) | 89 |
| 340 | 16 | cartddg (ref2015) | 160 |
| 340 | 16 | cartddg (talaris2014) | 70 |
| 340 | 1 | relax | 16 |
| 600 | 1 | relax | 40 |
| 900 | 1 | relax | 65 |
| 120;17* | 40 | flexddg (talaris2014) | 67 |

### Case study 1 - prediction of changes in folding free energy upon mutations and comparison with experimental values from the ThermoMut database

To illustrate the performance of the *ref2015* energy function, we performed folding free energy calculations with both the *cartddg* (**Figure 2**) and the *cartddg2020* (**Figure S1**) protocols and compared them to experimentally determined unfolding ΔΔG values. The following section illustrates, as an example, our findings when using the *cartddg* protocol. We downloaded the entire ThermoMut database ^26^ (ThermoMutDB) and selected four proteins as detailed in the Methods. In particular, we selected two bacterial enzymes with respectively 117 and 597 mutations, i.e., Enterobacteria phage T4 Endolysin, ENLYS (UniProt ID: P00720), and Staphylococcus aureus Thermonuclease, NUC (P00644). In addition, we performed the calculations on two human proteins of interest in health and disease, i.e., TP53 (P04637) and FKBP1A (P62942) with respectively 45 and 68 mutations with structural coverage. The secondary structure definition of the proteins was obtained from PDBe ^27^, and each position was annotated as either ɑ-helix, β-sheet, or loop in the wild-type structures. This case study aims at investigating the relationship between experimental and predicted values, per-mutation, when the data from all four proteins are pooled, allowing us to achieve better statistical power than considering each protein separately.

**Figure 2.**
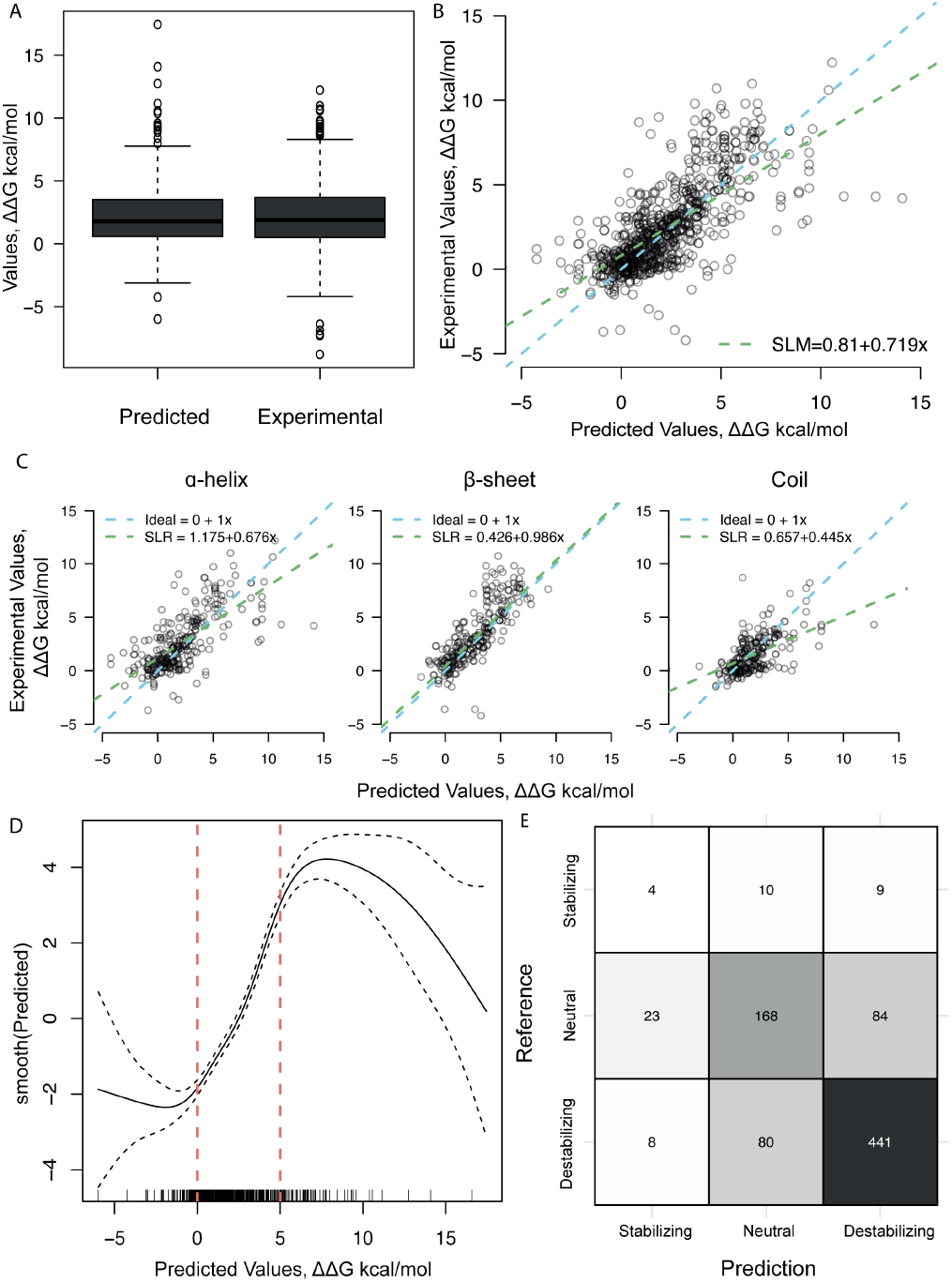
Comparison of changes in structural stability predicted with the *ref2015 cartddg* protocol and experiments. (A) Distribution of the predicted and experimental stability changes in kcal/mol. (B) Scatterplot of the ΔΔG values predicted by the *ref2015 cartddg* protocol and experimental values for the corresponding mutations. The blue line indicates a perfect correspondence between the variables. The green line is the fitted simple linear model. The model has an intercept of0.81, a slopeof 0.72, a variance (σ^2^) of 3.95, and a R^2^ of 0.44, a Pearson’s Correlation Coefficient of0.66, and a Mean Absolute Error between the predicted and experimental ΔΔGs (MAE) of1.39. (C) Scatterplots dividing the data by the wild-type secondary structure of the mutated position. The blue line indicates a perfect correspondence between the variables for each plot. The green line is the fitted simple linear model. Here, it is evident how the structured sections have a better correlation when compared to coils. This is likely due to the flexibility of the unstructured sections. ɑ-helices: Pearson’s correlation coefficient=0.70, slope=0.68, σ^2^=3.63, R^2^=0.49, MAE=1.53. β-sheets: Pearson’s correlation coefficient=0.69, slope=0.99, σ^2^=4.98, R^2^=0.49, MAE=1.38. Coil: Pearson’s correlation coefficient=0.57, slope=0.45, σ^2^=1.99, R^2^=0.33, MAE=1.22. (D) Generalized additive model (GAM) modeling the response variable, the experimental ΔΔG value, to a predictive variable, the predicted ΔΔG value, by estimating a smooth function, smooth (Predict). The smooth function has an effective degree of freedom of 6.5, quantifying the complexity of the line. The confidence interval is sufficiently narrow in the ΔΔG interval 0-5 kcal/mol to indicate that a linear relationship is present in this interval. (E) Confusion matrix where the experimental values are annotated as the reference values. The threshold used to define the classes is a ΔΔG of < −1 kcal/mol for stabilizing mutations, −1 < ΔΔG < 1 kcal/mol for neutral mutations and ΔΔG > 1 kcal/mol for destabilizing mutations. The resulting accuracy is 0.74.

To understand the agreement between the experimentally determined and the predicted stability, we performed a preliminary data exploration. Interestingly, data points from the experimental and prediction dataset are similarly distributed (**Figure 2A**), as corroborated by the Kolmogorov-Smirnov test (p=0.21).

We then tried to investigate the relationship between predicted and experimental data using a simple linear regression model, with the assumption that a perfect agreement between the experimental and predicted values would have an intercept of 0 and a coefficient of 1. The regression line has an intercept of 0.81 and a slope of 0.719 (**Figure 2B**). The variance of the linear model (σ^2^) is 3.95, and the model produces an R^2^ of 0.44, a Pearson correlation coefficient of 0.66, and a mean absolute error between the predicted and experimental ΔΔGs (MAE) of 1.39. The residuals plot for this model shows how the poor R^2^ value is at least partially due to systematic bias (**Figure S2**). This illustrates that a linear model does not completely explain the variance in the data we observed. To better understand this behavior, we tried to fit the data using a Generalized Additive Model (GAM) (**Figure 2D**). The resulting model has a roughly linear behavior in the ∼0-5 kcal/mol range but becomes less so at lower or larger ΔΔG values. Similarly, the confidence interval is very narrow in the linear regime interval, and it is wider for larger and smaller ΔΔG values, for which we have fewer data points. This observation is in alignment with Høie et al. ^28^, who found that ΔΔG predictions made with *ref2015* and the *cartesian2020* protocol in 29 proteins correlated with altered protein functions for ΔΔG > 4.5 kcal/mol, but the severity of the impact did not increase remarkably beyond this point. We then assessed the impact of the secondary structure on the performance of the prediction by building a simple linear model for each of the secondary structure groups, divided into ɑ-helices, β-sheets, and coil (**Figure 2C**). In fact, residues involved in structured regions are more likely to be part of the protein core, less flexible, and more sensitive to mutation with respect to solvent-accessible unstructured loops. The data points in the ɑ-helices and β-sheets are reasonably well correlated with Pearson correlation coefficients of 0.70 (slope=0.68, σ^2^=3.63, R^2^=0.49, MAE between predictions and experiments = 1.53) and 0.69, respectively (slope=0.99, σ^2^=4.98, R^2^=0.49, MAE between predictions and experiments = 1.38), while the correlation of the loop residues is 0.57 (slope=0.45, σ^2^=1.99, R^2^=0.33, MAE between predictions and experiments = 1.22), illustrating that the prediction is less consistent for unstructured regions. In this dataset, we noticed several outliers in which amino acid substitutions are predicted to have a large destabilizing impact, while the experiments find the variant to be neutral or mildly destabilizing. The experimental findings mostly align with the expectation that substitutions in flexible loops have mild effects on stability, although some loop substitutions may extend or create secondary structure elements, for example, as a result of substitutions from proline ^29^. The difference witnessed in this dataset is likely due to the fact that Rosetta allows some local main chain flexibility which may not be enough to represent the conformational variability that disordered regions experience in solution. We noticed similar behavior in applying FoldX, which we could mitigate with the usage of ensembles of structures generated, for example, by molecular dynamics simulations ^21,23,30^. It should be noted that the ɑ-helix mutations dataset also contains outliers. This dataset, however, has an overall better correlation with the experimental dataset, and the coefficient of its regression line is closer to 1. This suggests that changes in loops are more difficult to predict.

We then checked how good the performance of the predictions was when classifying mutations into destabilizing, neutral, and stabilizing. We did so by classifying all mutations causing stability changes above 1 kcal/mol as destabilizing, all mutations causing stability changes between 1 kcal/mol and −1 kcal/mol as neutral, and all mutations causing stability changes below −1 kcal/mol as stabilizing ^8,16^ and constructing a confusion matrix (**Figure 2E**). This confusion matrix yields an accuracy of 0.74. Where the accuracy in prediction is best for the destabilizing class (0.76), with high sensitivity (0.83), while the accuracy of the stabilizing class is only 0.56, with a sensitivity of 0.17, indicating that the destabilizing class is more likely to be correctly identified as compared to the stabilizing class. The neutral class has a similar performance to the destabilizing class (**Table S1**). In conclusion, this case study shows a good linear correlation between predicted and experimental values, especially in the 0-5 kcal/mol range; for larger and smaller values, while the trend is generally conserved, the relationship between the predicted and experimental values is less direct. We also show how predictions are more reliable for structured regions of the protein while correlation values are lower for unstructured regions. We performed the same analyses on the dataset obtained using the *cartesian2020* protocol, which showed similar trends overall (**Figure S1**).

### Case study 2 – prediction of changes in binding free energy for protein-short linear motifs (SLiMs) interactions

Within the protein-protein interaction landscape, intrinsically disordered proteins, or regions (IDPs, i.e., proteins that lack a defined tertiary structure or IDRs) have been proved to play an essential role in different biological events. IDP and IDRs include functional motifs known as Short Linear Motifs (SLiMs) that are important for the binding between IDPs and their target proteins ^31–33^. An example is the LC3 Interacting Region (LIR), i.e., a class of SLiMs involved in selective autophagy ^34^. One of the main features for regulating LIR binding to proteins of the LC3 family is through post-translational modifications (PTMs), especially through phosphorylation ^34^.

Here, we aim to show an application of the *flexddg* protocol to capture the changes in binding free energy upon phosphorylation or mutations in the core region of LIR-containing proteins.

First, we selected two examples of experimentally characterized phospho-regulated LIRs for which the structures were available on the Protein Data Bank, i.e., FUNDC1 in complex with LC3B (PDB entry, 2N9X ^35^) and PIK3C3 in complex with GABARAP (PDB entry, 6HOG ^36^). Experimental data on these two complexes through Isothermal Titration Calorimetry (ITC) and peptide arrays are available for these complexes, including the effects of phosphorylations or phospho-mimicking mutations ^35–37^. We applied the *flexddg* protocol with the *talaris2014* Rosetta energy function to investigate the effects of single and multiple phospho-mimetic mutations at the known phosphosites (see the Material and Methods section) since Rosetta does not currently provide parameters for phosphorylated residues. The results are described in detail below and reported in **Figure 3**.

**Figure 3.**
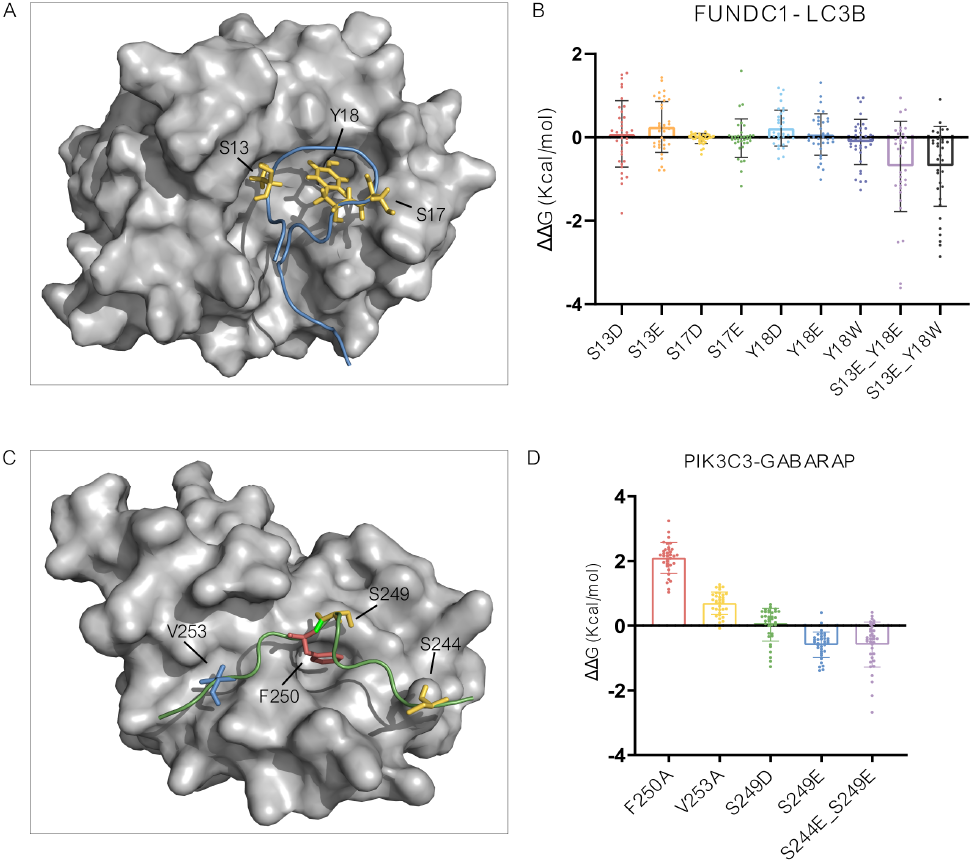
Prediction of changes in binding free energy using the *flexddg* protocol for protein interactions mediated by Short Linear Motifs. (A) FUNDC1 LIR peptide (blue) in complex with LC3B (gray) in the structure associated with the PDB entry 2N9X. The S13, Y18, and S17 phosphosites are shown as sticks and colored in yellow. (B) We report the predicted binding ΔΔGs for the single and double phosphomimetic mutations for the FUNDC1 LIR phosphosites for which experimental data are available for comparison. (C) PIK3C3 LIR peptide (blue) in complex with GABARAP (gray) in the structure associated with the PDB entry 6HOG. The S244 and S249 phosphosites are shown as sticks and colored in yellow, while the residues for binding to the GABARAP HP1 and HP2 pockets are shown as red and blue sticks, respectively. (D) We report the predicted binding ΔΔGs for single and double phosphomimetic mutations, along with mutations to alanine in the core motif of the PIK3C3 LIR, for which experimental data are available for comparison.

FUNDC1 is a mitophagy receptor that mediates the selective removal of damaged or superfluous mitochondria. It contains a canonical LIR (core region, 18-YEVL-21), which is necessary for the interaction with LC3 and, thus, for its role in mitophagy ^35^. FUNDC1 presents three experimentally validated phosphosites in or in the surroundings of its LIR motif: S13, S17, and Y18 (**Figure 3A**). Isothermal Titration Calorimetry (ITC) experiments with different FUNDC1 LIR peptides and LC3B reported a Kd of 0.40 ± 0.06μM for the wild-type variant. Phosphorylation at the S13 site resulted only in a slight decrease of the LC3B affinity (Kd= 0.60 ± 0.05μM) with respect to the wild type, while the phosphorylation of the Y18 site caused an approximately 5-fold increase in the Kd of interaction with LC3B (Kd= 1.72 ± 0.30μM). This increase is slightly augmented if both phosphorylations are combined (Kd= 2.00 ± 0.37μM) ^35^. Additionally, another work reported that phosphorylation of the S17 site increases the binding affinity with LC3B by approximately three folds ^37^. The *flexddg* protocol predicted the S13D and S13E substitutions to have neutral effects on the binding, in agreement with the experimental results (i.e., average ΔΔG < 0.25 kcal/mol).

However, the average ΔΔGs for the S17E and S17D mutations are also low, suggesting that, in this other case, the prediction cannot capture the changes in the binding affinity observed experimentally (**Figure 3A**). This is also the case for the phospho-mimetic mutations at the site Y18, whereas the combined effect of phospho-mimetic mutations at S13 and Y18 sites resulted in negative ΔΔG values, suggesting a stabilizing effect as observed experimentally (**Figure 3B**). Nevertheless, we also observed that the standard deviation is very high for these predictions, not allowing for quantitative conclusions.

The second protein of interest, PIK3C3, is a class III phosphoinositide 3-kinase enzyme of the PtdIns3K complexes (class III phosphatydylinositol 3-kinase complexes I and II) involved in the initiation of autophagy.

PIK3C3 presents a canonical F-type LIR (250-FELV-253) required for the interaction with GABARAP and GABARAPL1^36^. The effect of phosphorylations at S244 and S249 was assessed with ITC. In these experiments, the substitution of both the phosphosites with glutamate caused a 17-fold increase in GABARAP binding compared to the wild-type variant (Kd= 49.5 ± 3.9 μM). Moreover, peptide array experiments showed an increase in the binding affinity of the LIR peptide with all the LC3 family members for the phosphomimetic S249E variant ^36^.

To assess the potential of the *flexddg* protocol in capturing the effects induced by the phosphorylations of the PIK3C3 LIR, we modeled the S249E variant and a variant including phosphomimetic mutations at both the S44 and S249 sites (i.e., S244E_S249E, **Figure 3C**). We also tested the effect of the S249D substitution as a possible phosphomimetic variant, even if no experimental data are available for this mutation. We can observe that using S249D as a phosphomimetic variant does not provide the same result as introducing a glutamate (**Figure 3D**). This supports the notion that aspartate and glutamate cannot be always used as phosphomimetics in an interchangeable manner. The S249E variant seems to have a slightly stabilizing effect on the binding (average ΔΔG= −0.59 kcal/mol) and, in general, values of ΔΔG lower than 0 across the 35 independent runs (**Figure 3D**). This is in partial agreement with the peptide array results mentioned above. The double mutant variant S244E_S249E does not seem to increase the binding to the extent expected from the ITC data, suggesting that the *flexddg* protocol cannot be widely used to address multiple amino acid substitution. Indeed, the ΔΔG values predicted for the S244E_S249E variant are similar to the ones of the single amino acid substitutions.

Furthermore, we evaluated if the *flexddg* protocol could provide insights on the effects of mutations in SLiMs where PTMs are not involved. In the case of LIRs, the interaction between an LIR-containing protein and an LC3 family member is mainly driven by two residues of the LIR motif, which are in position 1 and 4 of the core motif and that bind to the Hydrophobic Pocket 1 (HP1) and the Hydrophobic pocket 2 (HP2) residues of the LC3 protein, respectively ^34^. Thus, we tested the capability of the *flexddg* protocol with the *talaris2014* energy function to predict the impact of the known detrimental mutations F250A (residue for interaction with HP1 pocket) and V253A (HP2 pocket) of PIK3C in complex with GABARAP (PDB entry, 6HOG ^38^). As expected, the *flexddg* protocol could predict the destabilizing effect of these mutations on the binding with GABARAP with predicted average ΔΔG values of 2.096 kcal/mol for F250A and 0.695 kcal/mol for V253A (**Figure 3D**). The main effect is triggered by the mutation of the residue for the binding to the HP1 pocket, in agreement with what is known from structural studies on LIR-LC3 protein interactions ^34^.

Overall, these applications illustrate the potential and some of the limitations of the RosettaDDGPrediction workflow, where two main challenges are the prediction of effects that are related to increased binding affinity (i.e., stabilizing mutations) and the reliable prediction of effects induced by phosphorylation with the usage of phosphomimetic mutations only.

### Case study 3 – influence of the source of initial structures for the calculations

The advent of AlphaFold has revolutionized molecular modeling and structural biology ^39^, resulting in models of 3D structures of proteins with resolutions comparable to those achievable with experimental approaches. Currently, the AlphaFold database contains over 360.000 predicted protein structures of 21 model organisms ^40^, providing a rich source of structures for in silico mutational scans as the ones that can be performed by MutateX or RosettaDDGPrediction. Here, we carried out an investigation to evaluate the influence of using a model based on AlphaFold2 with respect to a good resolution X-ray structure of the same protein. For this goal, we used as a case study the DNA binding domain (DBD) of p53, for which experimental data are also available on 31 mutant variants from ThermoMutDB ^26^. We evaluated the agreement between our calculated and experimentally available data, using the same parameters and energy functions, either the *cartddg* or *cartddg2020* protocol, and the two different starting structures. We have also included in the comparison a variant of *cartddg* in which we increased the numbers of runs per mutation up to ten, to determine whether it would improve our results. As the final ΔΔG depends on the values obtained by the single runs (even though it is calculated differently in the *cartddg* and *cartddg2020* protocols - see above), we expect that increasing the number of samples might lead to better converged final ΔΔG values if using just three runs is insufficient. We have measured the agreement through several metrics, such as the Pearson correlation coefficient, MAE, and a ROC curve.

We performed most of our comparison considering runs performed with the *cartddg* protocol. Therefore, in this section, we will be referring to them unless stated otherwise.

We obtained a similar pattern when comparing predictions and experiments using the experimental structure and the Alphafold2 model (**Figure 4A**) with a positive linear correlation, as quantified by Pearson’s correlation coefficient (**Figure 4B**). The highest Pearson’s correlation coefficient obtained was 0.79 using the scoring function *talaris2014* with the AlphaFold2 model and ten runs (**Figure 4B**), although all runs, including the ones using the *cartddg2020* protocol, achieved a correlation in the 0.57-0.79 range. Values ranging from 0.74 to 0.79 were obtained by all runs using the X-ray structure and by *talaris2014* with AlphaFold2 using three or ten runs. Using *ref2015* with the AlphaFold2 model led to a slightly worse correlation of 0.57, for three runs and 0.68 for ten runs. The runs with *ref2015* energy function and the *cartddg* protocol (X-ray structure) using ten runs, had the smallest Mean Absolute Error (MAE) of 0.90 kcal/mol **(Figure 4C**), meaning that it had the lowest average distance between predicted and target values among the different tested methods. It was closely followed by *talaris2014* with the AlphaFold2 structure using three and ten cycles with 0.92 and 0.91, respectively. The rest of the combinations had a MAE between 0.95 and 1.28.

**Figure 4.**
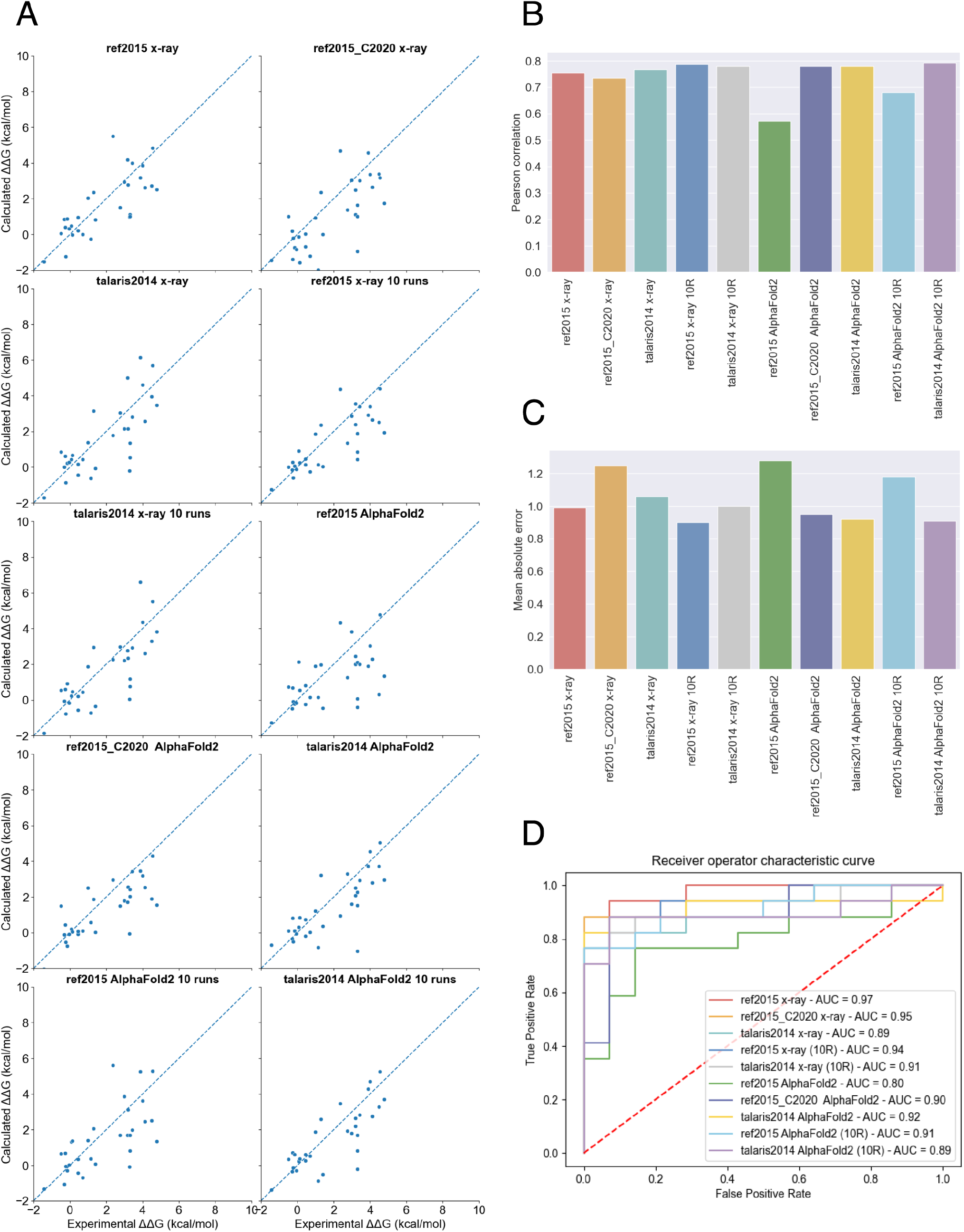
Comparison of experimental and predicted ΔΔGs using p53 as a case study. ΔΔG values were predicted using Rosetta version 3.12 withthe *ref2015* and *talaris2014* scoring functions, and the *cartddg* and *cartddg2020* protocols (referred to as “c2020” in the figure). We used the X-ray structure (PDB entry 2XWR) and a model from the AlphaFold2 database for the residues 91-289 of p53 as initial structures, using our default number of runs (three) or ten runs (these are referred to as “R10”). (A) Experimental vs. predicted ΔΔG values. (B) Pearson’s correlation coefficient between experimental and predicted values. (C) Mean Absolute Error (MAE) between experimental and predicted values. (D) Receiver operator characteristic (ROC) curve. The classification for this curve was done by considering the changes of free energy values reported in ThermoMutDB as ground truth, using 1.2 kcal/mol as ΔΔG cut-off to distinguish between destabilizing and non-destabilizing mutations (see Methods). The same criterion was used for the predicted mutations.

Considering the ROC curve, we considered experimental free energy changes from ThermoMutDB as ground truth and partitioned our dataset into destabilizing and non-destabilizing mutations depending on whether our prediction or ground truth had ΔΔG >= 1.2 kcal/mol. The best area under the curve (AUC) was achieved by using the scoring function *ref2015* using the *cartdgg* protocol and the X-ray structure, yielding a value of 0.97 (**Figure 4D**). In general, the different scoring functions and structures behaved similarly.

We did not appreciate great differences in the performance of our predictions when using the experimental X-ray structure or the AlphaFold2 model, with the only exception of using the *ref2015* energy function and the *cartddg* protocol with the AlphaFold2 model, which had a lower correlation and ROC AUC with respect to the other cases. Increasing the number of runs also slightly improved the performance, but with the trade-off of a considerably increased computing time. Finally, we obtained mixed results when comparing the *ref2015* X-ray three-cycles run performed using *cartddg* with the corresponding *cartddg2020* run. We did not see any appreciable improvement when using *cartddg2020* on the X-ray structure, as the *cartddg2020* run has a slightly lower correlation (0.74 vs. 0.76), higher MAE (1.25 vs. 0.99 kcal/mol), and lower AUC (0.95 vs. 0.97) considering the experimental data. Nonetheless, using *cartddg2020* with the AlphaFold2 model rescued the subpar performance of *ref2015* in this case, as all its performance measures are more similar to those of the other cases.

It should be noted that the DNA binding domain in the p53 AlphaFold2 model, ranging from residues 91 to 289, features a good per-residue confidence score (pLDDT) score, mostly above 70, meaning that more tests on models or regions with lower quality should be carried out to determine whether our findings can be generalized.

### Case study 4 – variants predisposing to childhood cancer

In a recent study, 198 samples from different childhood cancer types were analyzed with regard to germline variation and cancer predisposition ^41^. Among these, different variants of unknown significance (VUS) have been found with a frequency of < 1 % in the healthy population. Approximately 20% of the patients investigated had VUS in DNA repair pathway genes. In addition, we carried out new analyses on a larger dataset accounting for more than 500 germline samples from Danish children. The selection criteria for the proteins and the variants included in the study are described in detail in the Material and Methods and in **Figure S3**. We retained 14 proteins, i.e., ERCC4, BLM, FANCA, FANCE, FANCF, FANCG, FANCI, FANCL, MLH1, MSH2, MSH6, NBN, RAD51C, and RFWD3 for structure-based calculations of the changes in folding ΔΔGs for the VUS. All these genes are either classified as tumor suppressor genes in the COSMIC Cancer Gene Census v96 ^42^ or from literature for FANCI^43^ and RAD51C^44^.

Since mutations in tumor suppressor genes are generally causing loss-of-function in cancer^45^, we were interested in identifying VUS that have a destabilizing effect on the protein structure and thus result in positive predicted ΔΔG values upon mutation. These variants could be relevant to investigate further in terms of genomic alterations predisposing to cancer. To this aim, we retained the variants for which we had structural coverage with AlphaFold2 and high confidence scores (**Table S2**) for a total of 150 variants analyzed (**Figure 5-7**). According to searches in ClinVar^46,47^, some of the variants were already annotated as benign or likely benign but not related to childhood cancer. On the other hand, only T1131A in FANCA was found as pathogenic. The remaining were not deposited in ClinVar or annotated as unknown significance or with conflicting evidence, emphasizing the importance of additional analyses to understand the effects at the protein level.

**Figure 5.**
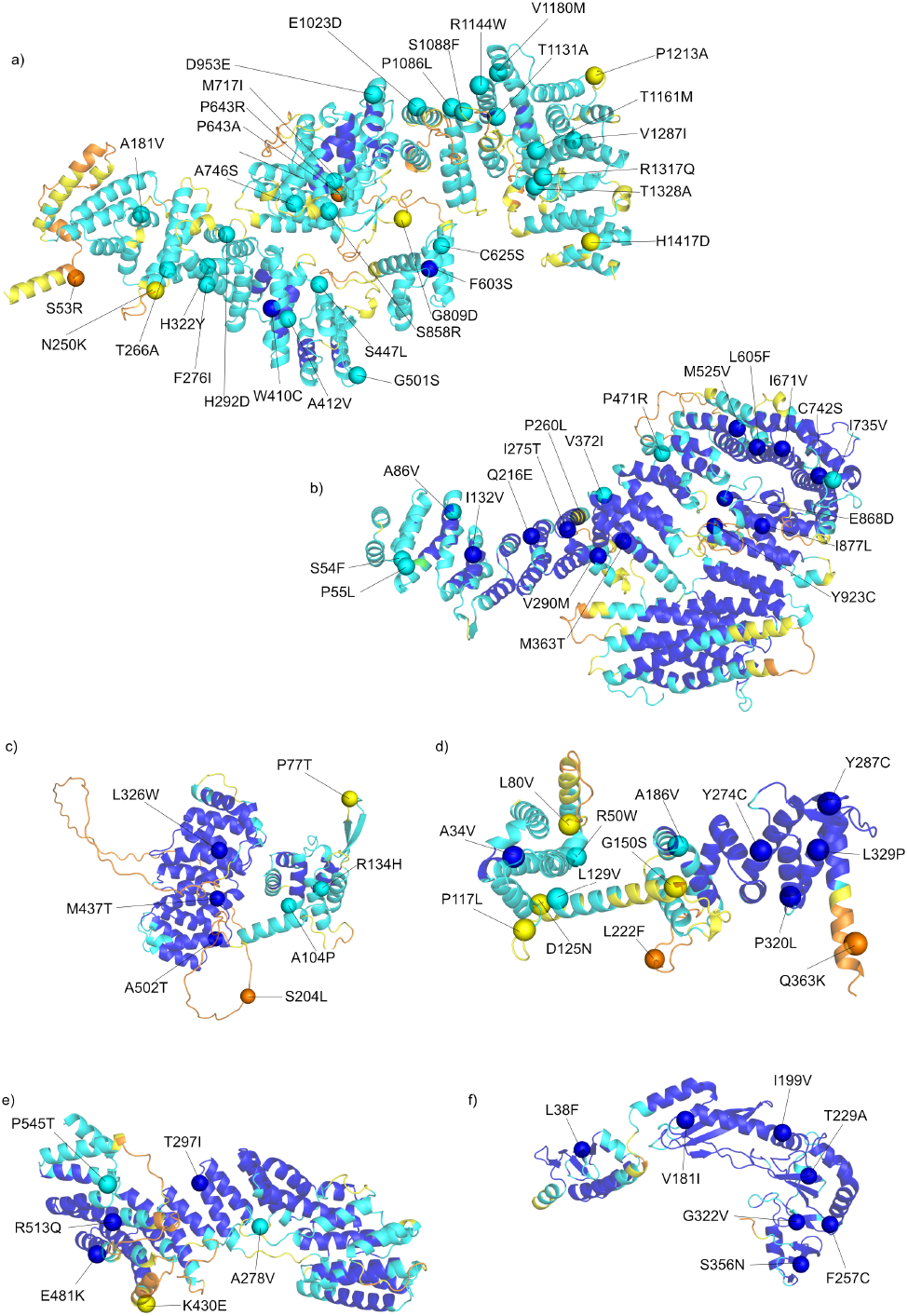
Trimmed AlphaFold structures of the FA (Fanconi Anemia) proteins selected for the case study 4. Cartoon representation of (A) FANCA_37-1441_, (B) FANCI_1-1279_, (C) FANCE_12-534_, (D) FANCF_2-369_, (E) FANCG_12-616_ and (F) FANCL_1-375_. The proteins are colored according to the AlphaFold2 pLDDT score: very low (yellow, pLDDT > 50), low (orange, 50 < pLDDT < 70), confident (light blue, 70 < pLDDT < 90), and very high (blue, pLDDT > 90). The Cɑ of the residues found mutated in pediatric cancer patients are shown as spheres and labeled.

**Figure 6.**
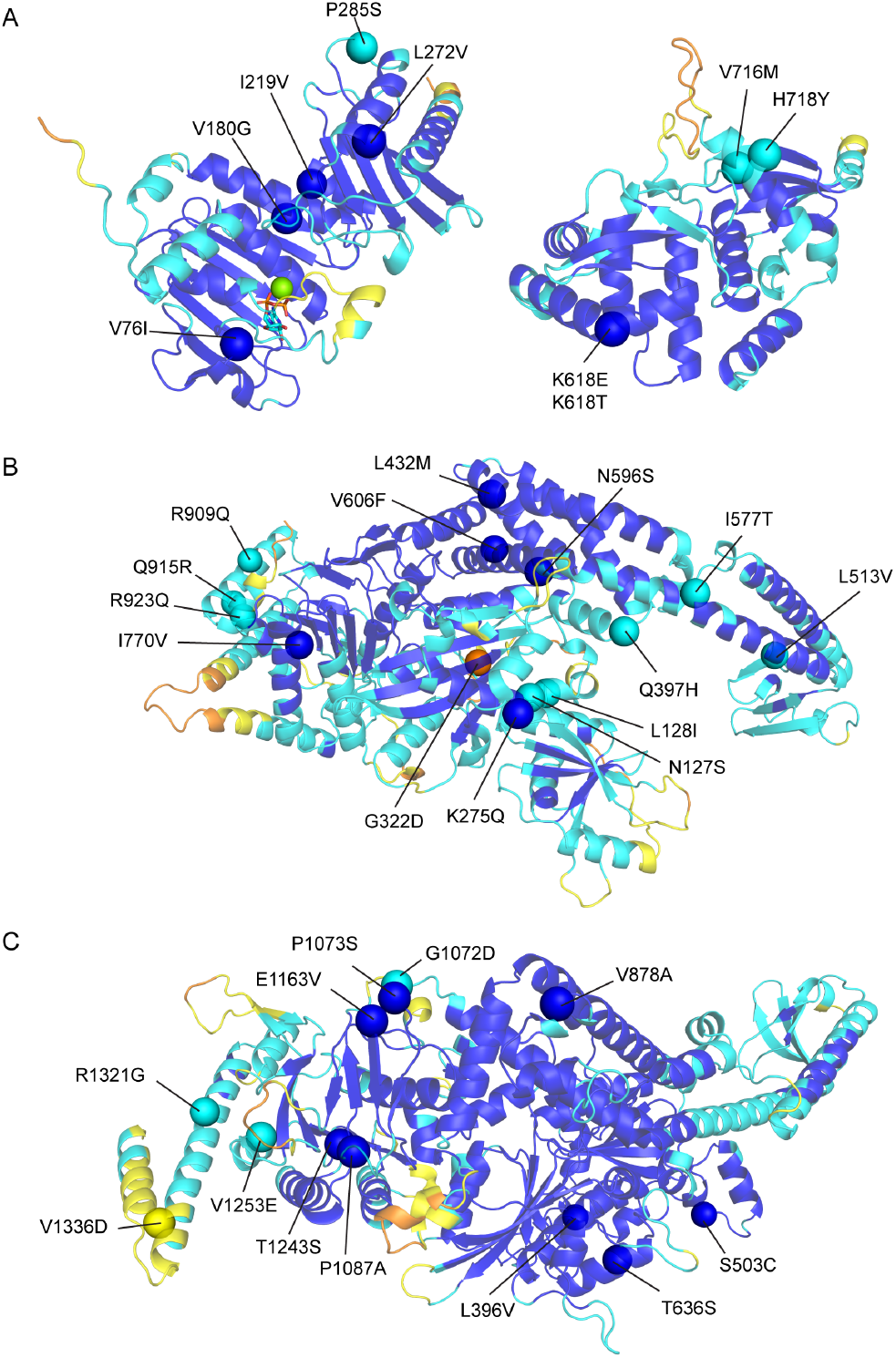
Trimmed AlphaFold structures of the of the DNA mismatch repair proteins selected for the case study 4. Cartoon representation of (A) MLH1_1-341_ and MLH1_501-756_, (B) MSH2_1-934_ and (C) MSH6_362-1360_. The proteins are colored according to the AlphaFold2 pLDDT score: very low (yellow, pLDDT > 50), low (orange, 50 < Plddt < 70), confident (light blue, 70 < pLDDT < 90), and very high (blue, pLDDT > 90). The Cɑ of the residues found mutated in pediatric cancer patients are shown as spheres and labeled.

**Figure 7.**
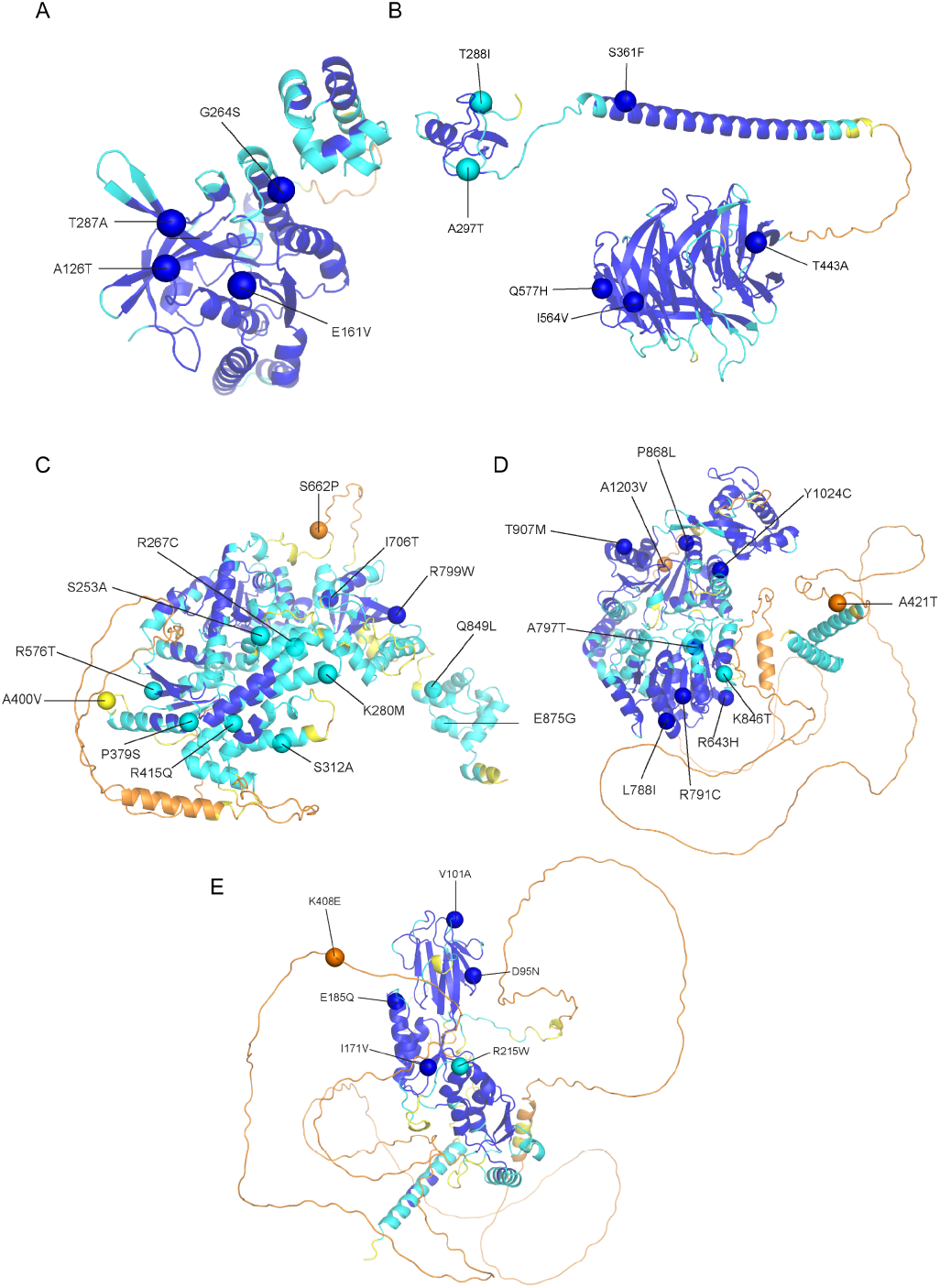
Trimmed AlphaFold structures of the proteins promoting the double-strand break (DBS) repair (RAD51C, RFWD3, ERCC4, NBN) and RECQ helicase (BLM) selected for proteins used for the case study 4. Cartoon representation of (A) RAD51C_13-350_ (B) RFWD3_284-774_ (C) ERCC4_12-914_ (D) BLM_368-1290_ and (E) NBN_1-749_. The proteins are colored according to the AlphaFold2 pLDDT score: very low (yellow, pLDDT > 50), low (orange, 50 < pLDDT < 70), confident (light blue, 70 < pLDDT < 90), and very high (blue, pLDDT > 90). The Cɑ of the residues found mutated in pediatric cancer patients are shown as spheres and labeled.

In this example, we applied the *cartddg2020* protocol, which considers the ΔΔG value referring to the mutant structure with the lowest total energy. We retained, as predicted destabilizing, the variants with ΔΔG values > 1 kcal/mol (see Methods) and confirmed destabilizing by calculations with MutateX (**Table S3, Table 2**). Indeed, the *foldX5* energy function, which is applied in the MutateX protocol, is effective in capturing loss-of-function mutations^48^. Of note, the pathogenic variant T1131A is not predicted to destabilize the structure of FANCA by both Rosetta and FoldX calculations (**Table S3**). We hypothesize that the detrimental effects triggered by this variant could be due to other properties such as impaired activity, interactions, or post-translational modifications at the cellular level. Experimental studies at the cellular level confirm that the T1131A substitution does not affect the protein levels, in agreement with a neutral effect on the folding ΔΔGs ^49^ and that the phenotype reflects a functional impairment that has a mild impact on MMC drug sensitivity and the monoubiquitinating of another protein ^49,50^. T1131A could be further investigated using our recently proposed multi-layered structural framework for variant annotations in proteins ^20,21,51^.

**Table 2.**
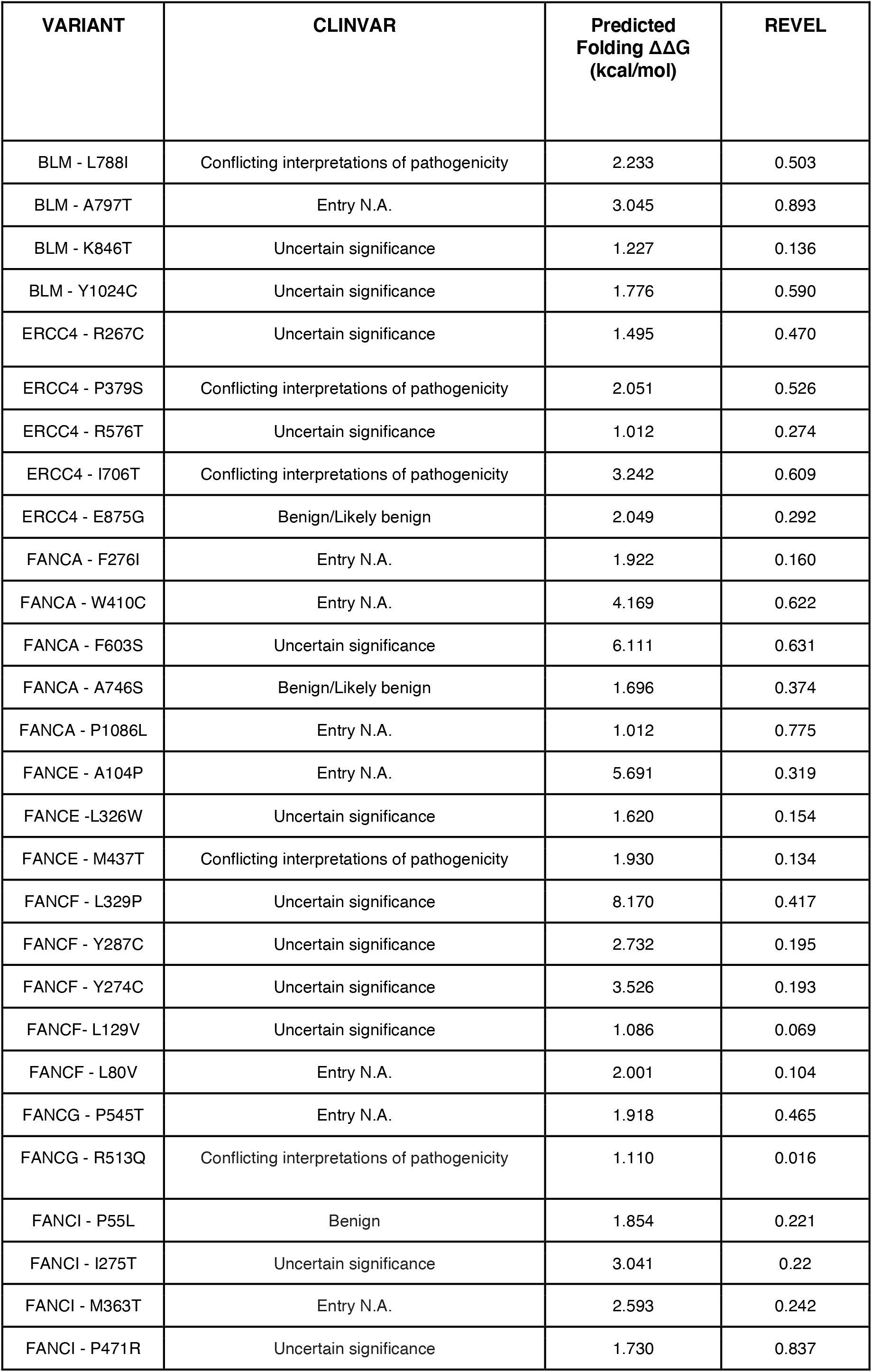
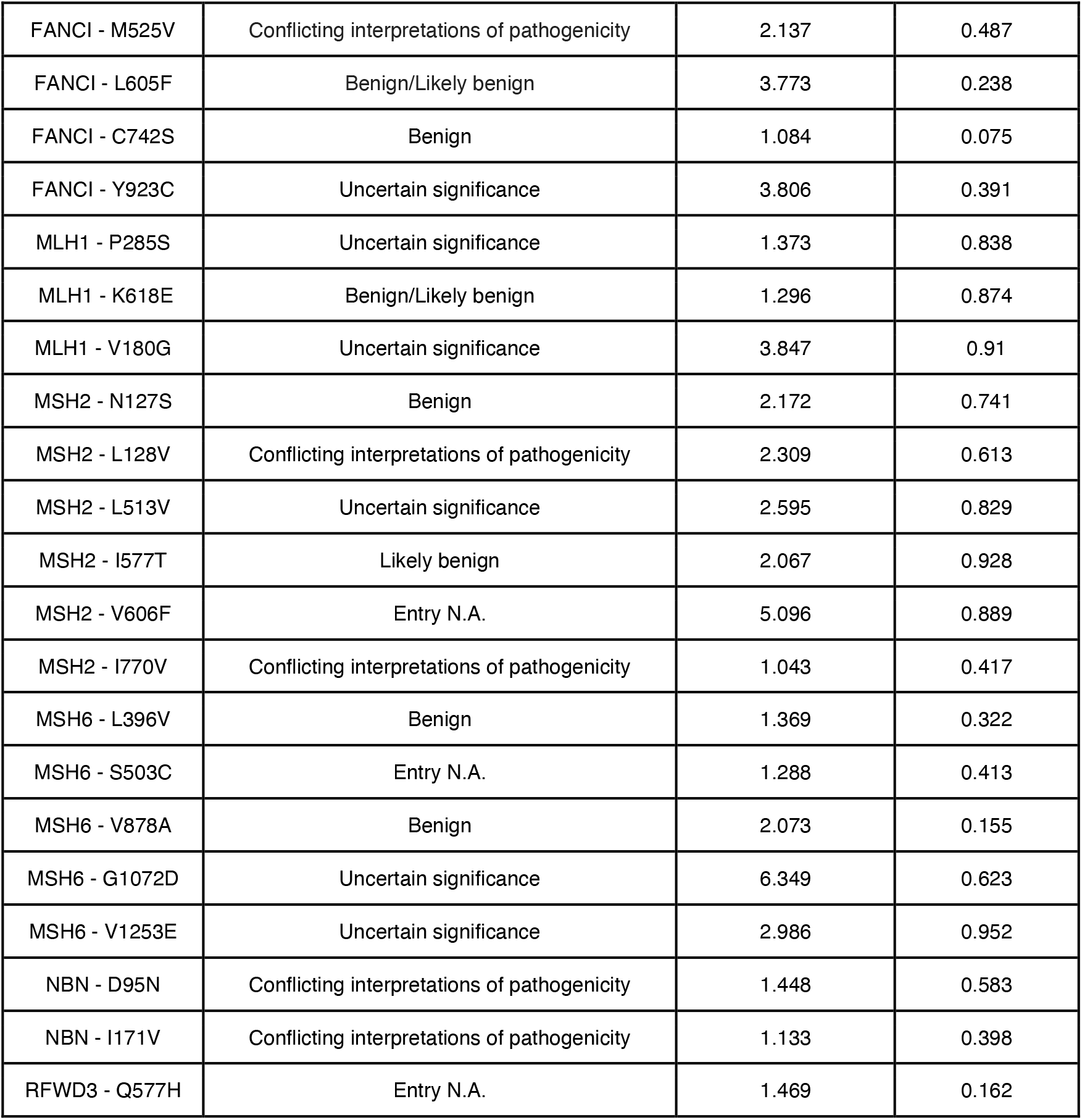
Summary of predicted ΔΔGs for Variants of Unknown Significance in childhood cancer. ‘NA’ indicates ‘Not Available’. We did not report RAD51C and FANCL in the table since all the variants analyzed here for these proteins were predicted with neutral effects for stability.

| VARIANT | CLINVAR | Predicted Folding $\Delta\Delta G$ (kcal/mol) | REVEL |
| --- | --- | --- | --- |
| BLM - L788I | Conflicting interpretations of pathogenicity | 2.233 | 0.503 |
| BLM - A797T | Entry N.A. | 3.045 | 0.893 |
| BLM - K846T | Uncertain significance | 1.227 | 0.136 |
| BLM - Y1024C | Uncertain significance | 1.776 | 0.590 |
| ERCC4 - R267C | Uncertain significance | 1.495 | 0.470 |
| ERCC4 - P379S | Conflicting interpretations of pathogenicity | 2.051 | 0.526 |
| ERCC4 - R576T | Uncertain significance | 1.012 | 0.274 |
| ERCC4 - I706T | Conflicting interpretations of pathogenicity | 3.242 | 0.609 |
| ERCC4 - E875G | Benign/Likely benign | 2.049 | 0.292 |
| FANCA - F276I | Entry N.A. | 1.922 | 0.160 |
| FANCA - W410C | Entry N.A. | 4.169 | 0.622 |
| FANCA - F603S | Uncertain significance | 6.111 | 0.631 |
| FANCA - A746S | Benign/Likely benign | 1.696 | 0.374 |
| FANCA - P1086L | Entry N.A. | 1.012 | 0.775 |
| FANCE - A104P | Entry N.A. | 5.691 | 0.319 |
| FANCE - L326W | Uncertain significance | 1.620 | 0.154 |
| FANCE - M437T | Conflicting interpretations of pathogenicity | 1.930 | 0.134 |
| FANCF - L329P | Uncertain significance | 8.170 | 0.417 |
| FANCF - Y287C | Uncertain significance | 2.732 | 0.195 |
| FANCF - Y274C | Uncertain significance | 3.526 | 0.193 |
| FANCF - L129V | Uncertain significance | 1.086 | 0.069 |
| FANCF - L80V | Entry N.A. | 2.001 | 0.104 |
| FANCG - P545T | Entry N.A. | 1.918 | 0.465 |
| FANCG - R513Q | Conflicting interpretations of pathogenicity | 1.110 | 0.016 |
| FANCI - P55L | Benign | 1.854 | 0.221 |
| FANCI - I275T | Uncertain significance | 3.041 | 0.22 |
| FANCI - M363T | Entry N.A. | 2.593 | 0.242 |
| FANCI - P471R | Uncertain significance | 1.730 | 0.837 |
| FANCI - M525V | Conflicting interpretations of pathogenicity | 2.137 | 0.487 |
| FANCI - L605F | Benign/Likely benign | 3.773 | 0.238 |
| FANCI - C742S | Benign | 1.084 | 0.075 |
| FANCI - Y923C | Uncertain significance | 3.806 | 0.391 |
| MLH1 - P285S | Uncertain significance | 1.373 | 0.838 |
| MLH1 - K618E | Benign/Likely benign | 1.296 | 0.874 |
| MLH1 - V180G | Uncertain significance | 3.847 | 0.91 |
| MSH2 - N127S | Benign | 2.172 | 0.741 |
| MSH2 - L128V | Conflicting interpretations of pathogenicity | 2.309 | 0.613 |
| MSH2 - L513V | Uncertain significance | 2.595 | 0.829 |
| MSH2 - I577T | Likely benign | 2.067 | 0.928 |
| MSH2 - V606F | Entry N.A. | 5.096 | 0.889 |
| MSH2 - I770V | Conflicting interpretations of pathogenicity | 1.043 | 0.417 |
| MSH6 - L396V | Benign | 1.369 | 0.322 |
| MSH6 - S503C | Entry N.A. | 1.288 | 0.413 |
| MSH6 - V878A | Benign | 2.073 | 0.155 |
| MSH6 - G1072D | Uncertain significance | 6.349 | 0.623 |
| MSH6 - V1253E | Uncertain significance | 2.986 | 0.952 |
| NBN - D95N | Conflicting interpretations of pathogenicity | 1.448 | 0.583 |
| NBN - I171V | Conflicting interpretations of pathogenicity | 1.133 | 0.398 |
| RFWD3 - Q577H | Entry N.A. | 1.469 | 0.162 |

We also observed that one variant annotated as benign in ClinVar (i.e., L605F FANCI) has predicted changes in folding ΔΔG higher than 3 kcal/mol and is, therefore, classified as destabilizing for the structural stability by our analysis. The variant has been characterized at the cellular level, showing decreased protein levels when compared to the wild-type, which confirms our prediction^52^. On the other hand, the variant P55L (predicted folding ΔΔG < 2.0 kcal/mol) was expressed at the same level as the wild-type. In addition, other benign variants according to ClinVar classification have been found in the range of 1-2 kcal/mol (**Table 2**). This observation suggests that variants for which the predicted changes in stability are within 1-3 kcal/mol should be further investigated to evaluate if they could result in neutral effects at the cellular level. In the case of MSH2, for example, it has been shown that a predicted destabilization of more than 3 kcal/mol is sufficient to cause cellular degradation of the protein ^53^.

According to the results in **Table2** and the observation above, if we consider folding ΔΔG values higher than 3 kcal/mol, our analyses suggest a number of VUS that could predispose to loss-of-function through destabilization of the protein structure and have a high REVEL score which further support their possible pathogenic impact (i.e., A797T in BLM, I706T in ERCC4, W410C and F603S in FANCA, L329P in FANCF, V180G in MLH1, V606F in MSH2, G1072D in MSH6). Of note, the mutation L329P in FANCE has been suggested to disrupt the stability of the catalytic module of the protein in a previous structural study ^54^.

## Discussion

We developed RosettaDDGPrediction moved by the need to provide easy and scalable access to Rosetta-based approaches to predict free energy changes in proteins upon mutations. RosettaDDGPrediction takes care of the whole process by performing a large number of ΔΔG predictions in an efficient and scalable manner, making a high-throughput calculations with Rosetta accessible, which is helpful for both extensive mutational scans and structured benchmarks.

RosettaDDGPrediction is, to our knowledge, the first wrapper devised to integrate state-of-the-art Rosetta-based protocols for the predictions of free energy changes upon mutation under a uniform framework.

Furthermore, the software checks the success of the runs, aggregates the data in CSV tables that are easy to mine and generates visual reports. As these steps are independent, the aggregation and visualization tools can be used on different datasets. In addition, we support additional output formats compatible with the MutateX plotting scheme ^23^. At the same time, raw or aggregated data can be easily manipulated externally. RosettaDDGPrediction also devotes particular attention to ensuring technical reproducibility by being controlled through configuration files. Further developments of RosettaDDGPrediction will focus on integrating its functionalities within MutateX, to provide a method-agostic container to perform and collect high-throughput mutational scans in a reproducible, automatized, and sustainable manner.

In this context, the performances of RosettaDDGPrediction and MutateX are only as good as those of the Rosetta- and FoldX-based methods that they incorporate. Indeed, Rosetta-based protocols implemented so far rely on different sampling methods to obtain models of the mutant variant structures and on scoring the resulting structures via knowledge-based energy functions to predict changes in the folding and binding free energy upon mutation ^16,55^. However, more rigorous strategies are available to predict both the effect of mutations on the folding free energy and the binding free energy^14,56–58^. For example, approaches leveraging enhanced sampling along reaction coordinates designed to study binding and unbinding events are available ^59–61^.

The time and computational resources needed by these methods still prevent their usage for investigations going beyond a few mutations. In these contexts, which include, for instance, saturation mutagenesis scans, Rosetta- and FoldX-based protocols represent a good trade-off between accuracy and speed.

Nevertheless, Rosetta still presents a challenge when non-canonical residue types are considered. Indeed, while most non-canonical amino acids are supported, mutations to phosphorylated residues cannot be performed in either protocol to predict free energy changes. For this reason, including strategies circumventing this issue would greatly expand the application of RosettaDDGPrediction.

Furthermore, a milestone in structural bioinformatics has been reached lately, with the release of AlphaFold2 and its outstanding performance in the CASP14 challenge ^39^. Originally developed to solve the long-standing protein folding problem, AlphaFold2 has already seen many spin-off studies to assess its potential ^62–66^. So far, evidence suggests that AlphaFold2 cannot effectively predict changes in folding free energy upon mutation ^67–69^. However, more studies are needed to explore this possibility fully.

Our wrappers have been devised to be inherently extensible. As stated above, a long-term perspective may include transforming them into a more general platform for structure-based methods to predict free energy changes upon mutation based on freely accessible, open-source software. This will also allow us to support other energy functions or schemes for free energy calculations.

The efforts of centralizing the development of software for in-silico deep mutational scans using free energy functions will help to move a step forward toward a unified framework for high-throughput structure-based calculations of free energy changes upon mutation.

## Methods

### Case Study 1

The ThermoMut database ^26^ was downloaded on April 22, 2022, as a JSON file. We then processed the database following four main steps: (i) For each reported protein, we retained only the entries including single mutations with an experimental value of ΔΔG discarding entries with multiple mutations with a combined ΔΔG; (ii) we reversed the sign of all the ΔΔG values to fit the sign provided by the outputs of RosettaDDGPrediction, (iii) we retained information on pH values and experimental methods as metadata and (iv) we removed protein entries for which less than ten mutations were reported. Upon processing, we identified 133 proteins. We then searched for three-dimensional structures available for each protein in the Protein Data Bank. In this step, we retained matches that covered at least one mutation of interest. We retained only protein structures in their free state (i.e., not in a complex with other interactors) for a total of 121 target proteins, effectively removing twelve proteins where no structure or free state was found. We then selected two enzymes that included a large number of amino acid substitutions with structural coverage (i.e., ENLYS and NUC as represented by the PDB structures 1P7S^70^ and 1EY0 ^71^, and two human proteins of interest in health and disease (p53 and FKB1A as represented by the PDB structures 2XWR ^72^ and 2PPN ^73^ as case studies for this work. All are used as simplistic monomeric structures and chosen based on the coverage, quality, and lack of interactors. The experimental values obtained in an acidic or alkaline experimental setting (pH < 6 and pH > 8) were excluded, as the *ref2015* Rosetta energy function (Cartesian space version) is simulating an environment at pH 7.

This leaves 845 observations across the four proteins for pH values 6, 7, and 8 and three methodologies, two chemically denaturant-induced protein unfolding experimental protocols, guanidine hydrochloride (GdnHCl), Urea Denaturation (Urea), and one thermal denaturation protocol (Thermal). We modeled the experimental and predicted values using a simple linear model, analyzed the contribution of secondary structures, and built a generalized additive model, thereby defining the limitations of the model. Furthermore, we constructed a confusion matrix based on the thresholds ΔΔG < −1 kcal/mol for the stabilizing group, −1 kcal/mol > ΔΔG < 1 kcal/mol for the neutral group, and ΔΔG > 1 kcal/mol for the destabilizing group. Calculations were carried out with Rosetta 3.12.

### Case Study 2

We started from the phospho-regulated LIRs reported in our previous review article ^34^ and other literature search, and, for each of them, we verified if a complex with one of the LC3/GABARAP family members was available to use as starting structure for the mutational scan. We retained for the analyses the following complexes: LC3B:FUNCD1 (PDB entry 2N9X ^35^) and GABARAP:PIK3C3 (PDB entry 6HOG ^38^).

We reconstructed missing coordinates in the structures using MODELLER version 10.1^74^.

We used the *flexddg* protocol, as implemented in RosettaDDGPrediction, with the *talaris2014* energy function and Rosetta 3.12. Rosetta Energy Units (REUs) were converted to kcal/mol with the conversion factors provided for this energy function ^16^. We modeled the phosphorylated residues using phosphomimetic mutations to aspartic acid and glutamic acid for each phosphosite, and included also tryptophan for phosphtyrosine to identify possible effects due to steric hindrance. In the calculations, we used 35 000 backrub trials and an absolute score threshold for minimization convergence of 1 REUs. We generated an ensemble of 35 structures for each mutant variant and calculated the average ΔΔGs and the standard deviation among the individual binding free energies.

### Case Study 3

We retrieved experimental ΔΔG values from point mutations of the p53 DNA-binding domain from the online database ThermoMutDB. Since ThermoMutDB stores ΔΔG_u_ values, they were converted to ΔΔG_f_ by changing the sign to make them easily comparable with Rosetta output values. A total of 31 mutations were selected, and when multiple experimental values were reported for the same variant, the average of their ΔΔG_f_ was used.

We used two different structures. The first one consists of the X-ray crystallography of the PDB entry 2XWR, with a resolution of 1.68 Å, which covers the DNA-binding domain from residues 91 to 289 and includes the Zinc ion. The water molecules were removed using PyMOL [http://www.pymol.org/pymol]. We also used the model from the AlphaFold2 database, which was trimmed to cover the same residues as the experimental X-ray structure, from 91 to 289. The missing zinc ion was added using PyMOL, identifying its coordinates by rigid body superimposition with the original structure. Before, we verified that the residues which coordinate the zinc ion (C176, H179, C238, C242) had a good alignment and similar rotamer position between the two structures.

For the ΔΔG predictions, we mostly used the *cartddg* protocol with the *ref2015* and *talaris2014* scoring functions, each with three and ten sampling runs and Rosetta 3.12. We also used the *cartddg2020* protocol on both structures, but only using the *ref2015* scoring function and three runs.

The performance was measured by Pearson’s correlation coefficient, mean absolute error, and area under the curve (AUC) of the ROC curve. For the ROC curve, we used a threshold of 1.2 kcal/mol for the ThermoMutDB averaged values, meaning that mutations associated with a free energy change higher than 1.2 kcal/mol were considered destabilizing, according to the threshold selection proposed for p53 in our previous study ^20^. In comparison, mutations associated with a free energy change lower than the threshold were classified as non-destabilizing. The mean absolute error was calculated using the following equation:

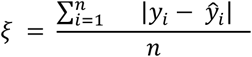

Here, y_i_ are the experimental values and *ŷ*_*l*_ ^ the predicted values.

### Case Study 4

We retrieved VUS in childhood cancer from a previous study ^41^. In addition, we analyzed a dataset including 566 samples from Danish children with different cancer types and Whole Genome Sequencing data. The sequencing data have been processed with a pipeline based on Sentieon using the build 38 (GATK resource bundle for hg38) as reference genome. Reads were aligned with BWA-MEM to the reference genome, and duplicate reads were removed. Reads were realigned around indels, and we applied Base Quality Score Recalibration together with the Haplotyper algorithm for variant calling (equivalent to the GATK Haplotype ^75^. Then, as suggested by GATK best practices, we used Variant Quality Score Recalibration, which is an advanced filtering technique used on the variant call set that models the technical profile of variants in a training set using machine learning and filters out potential artifacts from the callset. The filtered variants were uploaded to an in-house mySQL database where we linked them with information about genomic context (ENSEMBL v95), ENSEMBL consequences, deleteriousness-scores (CADD 1.6, REVEL, SIFT, PolyPhen) and variant frequency in the healthy population (GnomAD v3) based on their genomic position and alternate allele. In particular, we annotated the REVEL score ^76^ associated with the genomic mutation by using the publicly available dataset of precomputed scores, by matching genomic coordinates, annotated transcript for the mutation and alternate nucleotide. We could not annotate a REVEL score for four of the identified variants, most likely as they were not missense. Two of them caused early translation termination by introducing a stop codon in our reference BRCA2 transcript (13:g.32337185A>T and 13:g.32398489A>T, corresponding to p.Lys944* and p.Lys3326* at protein level).The other two (2:g.47607407G>A and 2:g.47607446G>A, corresponding to p.Arg923Gln and p.Gly936Asp in MSH2 at protein level for our reference transcript) were annotated as both missense and nonsense-mediated decay in our dataset, meaning they are annotated as nonsense-mediated decay for at least some of the MSH2 transcripts, and this is probably the reason they were not available in the REVEL database.

We retained, as VUS to investigate, those variants located in the coding regions and found with a frequency lower than 1% in GnomAD v3 (build 38) as a proxy for a healthy population. This threshold has been selected according to the guidelines for VUS studies ^77^. An illustration of the workflow for analyzing the sequencing data is provided in **Figure S2**.

We searched each variant in the selected 14 genes for the study in ClinVar ^46,47^ and retrieved annotations on them to verify if they are VUS, variants with conflicting evidence, or not reported yet in the database. To select the proteins and variants that can be investigated with RosettaDDGPrediction, we then searched in the AlphaFold2 database ^40^ for the corresponding protein structures and retained those that had structural coverage for the variants in regions with high confidence (pLDDT > 70). Cases in which the pLDDT score is low but located in loops that connect structured regions of folded domains were also retained for analyses. These regions are often very flexible in a protein structure, and it is thus expected that they could have a lower pLDDT score. The selected target proteins and corresponding variants are reported in **Table S2**. We analyzed 14 proteins and 132 variants in total.

We excluded mutations either not covered by our trimmed models or derived from an isoform different from the one available in the AlphaFold2 database. Concerning MSH2, we did not analyze G936D since our isoform had 934 residues, while R293Q refers to the A0A2R8YG02_HUMAN isoform ^78^. In the case of MSH6, T1125M was removed since derived from the A0A494C0M1_HUMAN transcript ^78^. Furthermore, the following seven variants found in FANCL were also disregarded: S356N, S356N, G322V, F257C, T229A, I199V, and V181I. These variants were generated from FANCL isoform 2 (ENST00000402135.8, Q9NW38-2, 380aa), which did not match the AlphaFold model for FANCL (ENST00000233741.9, Q9NW38, 375aa).

## Acknowledgements

Our research has been supported by Carlsberg Foundation Distinguished Fellowship (CF18-0314), Danmarks Grundforskningsfond (DNRF125), Hartmanns Fond (R241-A33877), LEO Foundation (LF17006) and NovoNordisk Fonden Bioscience and Basic Biomedicine (NNF20OC0065262).

This work is part of Interregional Childhood Oncology Precision Medicine Exploration (iCOPE), a cross-Oresund collaboration between University Hospital Copenhagen, Rigshospitalet, Lund University, Region Skåne, and Technical University Denmark (DTU), supported by the European Regional Development Fund. This work is also part of the Danish nation-wide research program Childhood Oncology Network Targeting Research, Organisation & Life expectancy (CONTROL) and supported by the Danish Cancer Society (R-257-A14720) and the Danish Childhood Cancer Foundation (2019-5934 and 2020-5769).

This paper was typeset with the bioRxiv word template by @Chrelli upon small modifications: www.github.com/chrelli/bioRxiv-word-template

## Author contributions

*Sora Valentina:* Conceptualization (RosettaDDGPrediction), Methodology (RosettaDDGPrediction), Software (RosettaDDGPrediction), Visualization (Figure 1), Writing - Original Draft Preparation (Abstract, Introduction, RosettaDDGPrediction, Discussion), Writing – Review & Editing

*Otamendi Laspiur Adrian:* Conceptualization (case study 4), Data Curation (case study 4), Formal Analysis (case study 4), Investigation (case study 4), Methodology (case study 4), Software (case study 4), Validation (case study 4), Visualization (Figure S3, Table S2, Figure 5, Table 2), Writing - Original Draft Preparation (case study 4), Writing – Review & Editing

*Degn Kristine*: Conceptualization (case study 1), Data Curation (case study 1), Formal Analysis (case study 1), Investigation (case study 1), Validation (case study 3), Visualization (Figure 2, Figure S1, Figure S2, Table S1), Writing - Original Draft Preparation (case study 1), Writing – Review & Editing

*Arnaudi Matteo*: Conceptualization (case study 2), Data Curation (cases study 2 and 4), Formal Analysis (cases study 2 and 4), Funding Acquisition; Investigation (cases study 2 and 4), Visualization (Figure 7), Writing - Original Draft Preparation (case study 2), Writing – Review & Editing

*Utichi Mattia*: Conceptualization (case study 2), Data Curation (case study 2), Formal Analysis (case study 2), Investigation (case study), Visualization (Figure 3), Writing - Original Draft Preparation (case study 2), Writing – Review & Editing

*Beltrame Ludovica*: Conceptualization (case study 4), Data Curation (case study 4), Formal Analysis (case study 4), Investigation (case study 4), Validation (case study 4), Visualization (Figure 6), Writing – Review & Editing

*De Menezes Dayana*: Conceptualization (case study 3), Data Curation (case study 3), Formal Analysis (case study 3), Investigation (case study 3), Visualization (Figure 4), Writing - Original Draft Preparation (case study 3), Writing – Review & Editing

*Orlandi Matteo*: Formal Analysis (case study 4), Writing – Review & Editing

*Rigina Olga*: Data Curation (case study 4), Software (case study 4), Writing – Review & Editing

*Wad Sackett Peter*: Data Curation (case study 4), Resources, Writing – Review & Editing

*Wadt Karin*: Data Curation (case study 4), Funding Acquisition, Resources, Supervision (case study 4), Writing – Review & Editing

*Schmiegelow Kjeld*: Funding Acquisition, Resources, Supervision (case study 4), Writing – Review & Editing

*Tiberti Matteo*: Conceptualization (case study 1), Supervision (case study 1 and 3), Validation (case study 1 and 3), Visualization (Figure 4), Writing - Original Draft Preparation (case study 1 and 3), Writing – Review & Editing

*Papaleo Elena*: Conceptualization (RosettaDDGPrediction and all case studies), Data Curation (case study 4), Formal Analysis (case study 4), Funding Acquisition, Project Administration, Resources, Supervision, Validation (case study 2 and 4), Visualization (Table 1, Table 2, Table S2), Writing - Original Draft Preparation (all Sections), Writing – Review & Editing

