## Supplementary Figure S1 for "RosettaDDGPrediction for high-throughput mutational scans: from stability to binding"

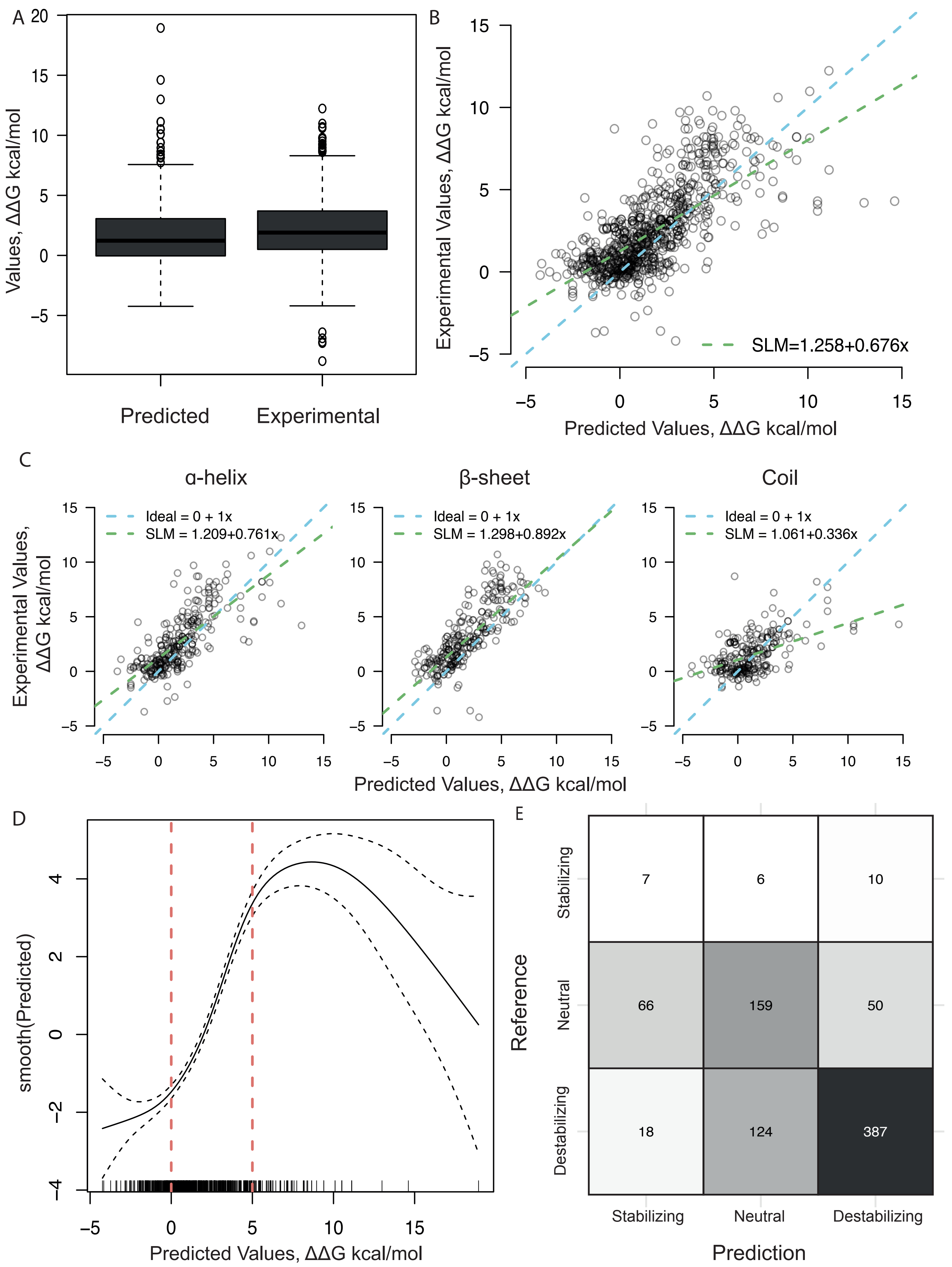


**Supplementary Figure S1.** Comparison of stability changes from the ref2015 cartesian2020 protocol and experimentally found values. (A) The distribution of the predicted and experimental stability changes in kcal/mol. (B) A scatterplot of the ΔΔG values predicted by the ref2015 cartesian2020 protocol and experimentally found values for the corresponding mutations. The green line is the fitted simple linear model. The model has an intercept=1.26, slope=0.68, σ2=4.06, R2=0.43, Pearson Correlation Coefficient=0.65, (C) Three scatterplots to illustrate the data as divided by the wild-type secondary structure of the mutated position. The green line is the fitted simple linear model. Here it is evident how the structured sections have a better correspondence as compared to the coils. α-helices: Pearson correlation coefficient=0.75, β-sheets: Pearson correlation coefficient=0.69, Coil: Pearson correlation coefficient=0.52, (D) A generalized additive model (GAM) modeling the response variable, experimental ΔΔG, to a predictive variable, predicted ΔΔG by estimating a smooth function, smooth(Predict). The smooth function has an effective degree of freedom of 5.5, quantifying the complexity of the line. The confidence interval is sufficiently narrow in the ΔΔG interval 0-5 kcal/mol to indicate that a linear relationship is present in this interval. (E) A confusion matrix where the experimental values are annotated as the reference values. The threshold used to define the classes is a ΔΔG of $< -1$ kcal/mol for stabilizing mutations, $-1 < \Delta\Delta G < 1$ kcal/mol for neutral mutations and $\Delta\Delta G > 1$ kcal/mol for destabilizing mutations. The resulting accuracy is 0.67.
