## Supplementary Figure S2 for "RosettaDDGPrediction for high-throughput mutational scans: from stability to binding"

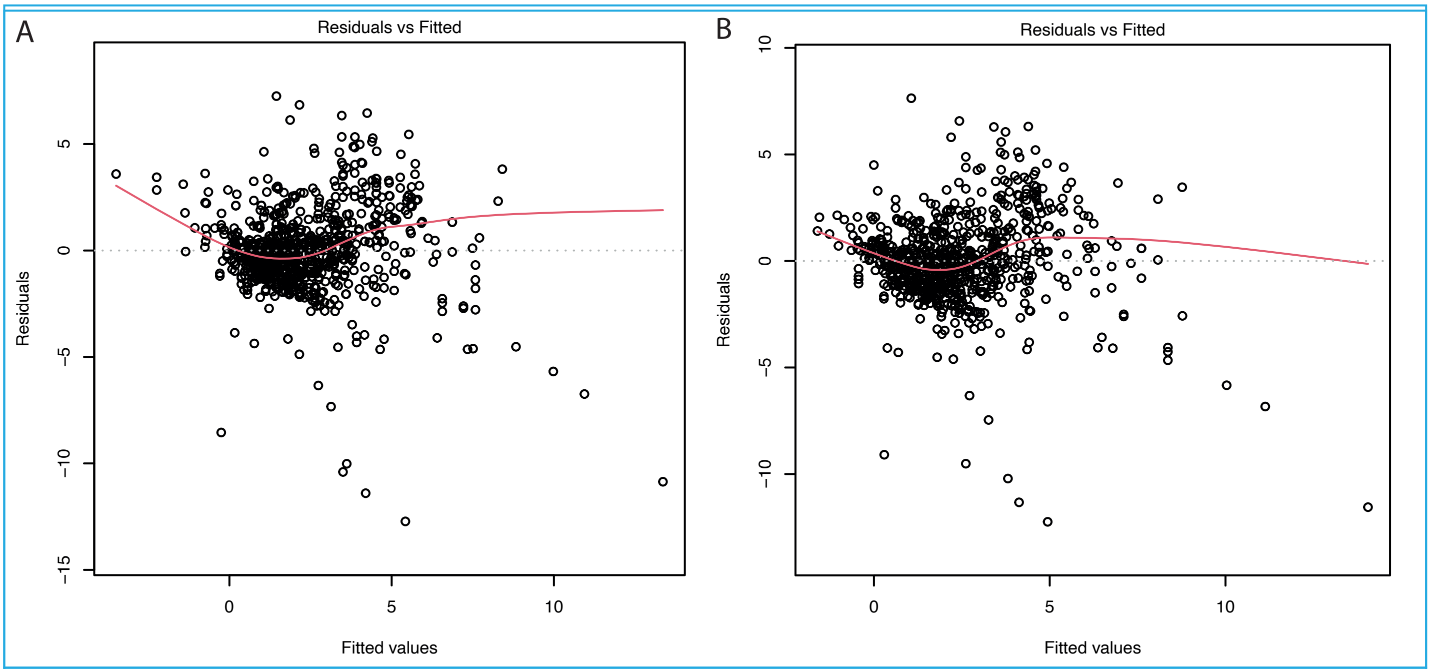


**Supplementary Figure S2.** Residual plots. (A) The difference between the observed and the fitted response values for the simple linear model from the ref2015 cartesian protocol. (B) The difference between the observed and the fitted response values for the simple linear model from the ref2015 cartesian2020 protocol. Both plots illustrate a biased fit, as they are not randomly scattered around the identity line y=0.
