## Supplementary Figure S3 for "RosettaDDGPrediction for high-throughput mutational scans: from stability to binding"

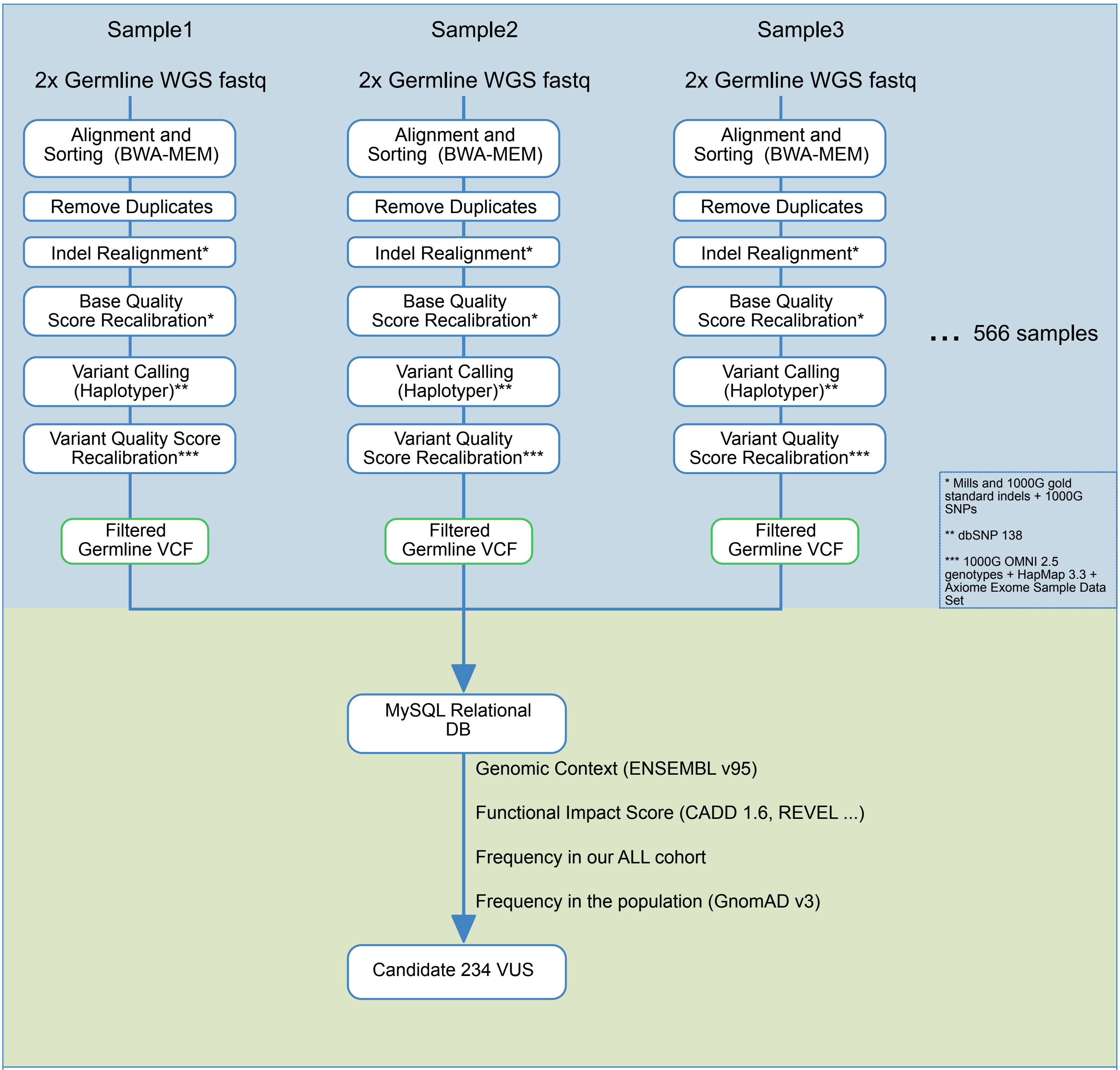


**Supplementary Figure S3**. Schematic representation of NGS bioinformatics pipeline for candidate VUS extraction and annotation. Fastq pairs for each sample (normal blood) are processed independently (n=566) with Sentieon® software based on GATK best practices. Reads are aligned to Human Genome build 38 (GATK resource bundle for hg38) with BWA-MEM algorithm. Duplicate reads are removed and reads are realigned around indels. After that, we perform Base Quality Score Recalibration to detect systematic errors produced by the sequencing machine. These last two steps are performed using as reference known SNPs and indels from Mills and 1000G projects (*). Then, variants are called with Haplotyper algorithm (equivalent to GATK Haplotype Caller (https://doi.org/10.1101/201178)) using SNPs from dbSNP 138 as known resource (**). Finally, Variant Quality Score Recalibration is performed which uses 1000G OMNI 2.5 genotypes, HapMap 3.3 genotypes and Axiom Exome Sample Data Set (***) to model the technical profile of the variants and filter our potential artifacts from our call set. Per sample call sets are independently uploaded into an in-house MySQL (MariaDB) database. For each variant, annotation for genomic context from Ensembl v95 is added, as well as Functional Impact scores (CADD 1.6, REVEL...) and frequency in the cohort and healthy population (GnomAD v3). Variants affecting the coding region with frequency in the healthy population of <1% were selected as candidate VUS (n=234).
